# Needle in a haystack: Fluorescence-guided 3D targeting for FIB milling in high-pressure-frozen samples

**DOI:** 10.64898/2026.11.20.768024

**Authors:** Oliwia Koczy, Fergus Tollervey, Zhengyi Yang, Anastasiia Babenko, Cristina Paulino, Simone Mattei

**Affiliations:** Molecular Systems Biology Unit, European Molecular Biology Laboratory, Heidelberg, Germany; Biochemistry Center Heidelberg, University of Heidelberg, Heidelberg, Germany; EMBL Imaging Centre, European Molecular Biology Laboratory, Heidelberg, Germany; Collaboration for joint PhD degree, Faculty of Biosciences, EMBL and Heidelberg University, Heidelberg, Germany

## Abstract

Rare subcellular structures are difficult to retain during lamella preparation for cryo-electron tomography, particularly in high-pressure-frozen specimens. Here we introduce two complementary workflows for three-dimensional fluorescence-guided milling on a commercial cryo-focused ion beam-scanning electron microscope with integrated fluorescence imaging. Iterative fluorescence correlation uses repeated rounds of imaging and milling to refine the target position, whereas bidirectional fluorescence correlation images a lifted-out block from two orthogonal directions. We validated both by targeting centrosomes in waffle-frozen early *Caenorhabditis elegans* embryos, each of which occupy approximately 0.007% of the embryo volume. We recovered recognizable centrosomes in 83% and 67% of lamellae examined by transmission electron microscopy, respectively, compared with 25% using a refined, 2D-guided high-throughput SOLIST workflow, run as a benchmark. Neither method requires hardware modification or fluorescent fiducial beads. These workflows provide accessible strategies for targeting rare subcellular structures during lamella preparation, with or without a lift-out hardware.

## INTRODUCTION

Cryo-focused ion beam (FIB) milling combined with cryo-electron tomography (cryo-ET) has enabled the structural analysis of macromolecular complexes directly within their native cellular environment^1–3^. Conventional *in situ* structural biology workflows rely on the analysis of thin samples that can be vitrified by plunge freezing and thinned by on-grid cryo-FIB milling^4^. The development of the first lift out approaches^3,5,6^ provided the opportunity of preparing thin lamellae also from thicker, multicellular specimens vitrified by high-pressure-freezing (HPF). However, the early methods were lengthy, relied on manual operations and were characterised by modest success rate. The broad adoption of lift out for *in situ* structural biology studies was enabled by the recent development of efficient and robust methods such as the serial lift-out^7^ and Serialized On-grid Lift-in Sectioning for Tomography (SOLIST)^8^, both of which have substantially tackled previous limitations in throughput, reliability, and user-friendliness. By enabling multiple lamellae to be generated from a single block of HPF material, these approaches have facilitated cryo-ET studies of specimens that are difficult or impossible to access using plunge freezing and conventional on-grid lamella preparation. These advances have enabled structural studies of extracellular matrices^9^, organoids and tissue biopsies^10–12^, plant roots^13^, and whole multicellular organisms^6,7^.

Locating a subcellular structure within HPF specimens remains difficult, as the exposed surface often provides only a few morphological landmarks for identifying an internal region of interest (ROI). Cryogenic correlative light and electron microscopy (cryo-CLEM) can identify fluorescently labelled regions and guide selection of a volume for milling or lift-out^14^. Such regional targeting is effective when the desired structures are abundant or occupy large volumes. For small, sparsely distributed targets, however, their position must be known in all three dimensions and maintained throughout preparation. A final lamella, only 200-400 nm thick, retains a small fraction of the starting volume, so even modest positioning errors can remove the target. Furthermore, accurate depth estimation is difficult with the optics used in most cryo-fluorescence microscopes. Air objectives, including those used in integrated cryo-FIB-scanning electron microscope (cryo-FIB/SEM) systems, limit numerical aperture and provide substantially poorer axial than lateral resolution^15–18^. In addition, the refractive-index mismatch between the vacuum or gas surrounding the sample and the vitreous ice distorts the apparent depth of fluorescent signals^19^. Cryo-immersion optics can improve optical performance^15,17,20,21^ but their use during milling is constrained by the vacuum environment and optical access requirements of a cryo-FIB-SEM chamber.

The limitations of ROI targeting by cryo-fluorescence imaging are compounded in thicker specimens. Whereas plunge frozen samples are typically below 10 µm thick, ‘waffle’ freezing typically produces samples approximately 25–35 µm thick^22^, and HPF specimens contained within metal planchettes range between 100 and 300 µm. With increasing depth, scattering reduces the fluorescence signal, and spherical aberrations caused by refractive-index mismatch and roughness of the ice surface broaden the point spread function^19,23^. Thus, correlative approaches developed for plunge frozen samples cannot always be applied directly to bulky specimens such as tissues and organoids^24^. Consequently, fluorescence imaging in lift-out studies has primarily been used to identify larger regions enriched in a fluorescent signal, thereby guiding selection of the volume to be extracted, rather than to precisely retain specific fluorescent puncta within the final lamella^7,10,12,25^.

Recent developments in correlative cryo-light and electron microscopy have enabled fluorescence information to guide FIB milling with increasing precision. Three-dimensional cryo-CLEM can register fluorescence volumes to FIB images using fiducial beads visible in both modalities^26^, and integrated fluorescence microscopes facilitate imaging during lamella preparation. However, introducing suitable beads into thick specimens and identifying them in electron or ion images can be difficult. Correlations must also accommodate drift and sample deformation as milling proceeds^27^. Coincident cryo-FLM-FIB-SEM configurations, in which fluorescence imaging can be performed during ion-beam milling, provide direct feedback on target position. However, the requirement for simultaneous optical, electron, and ion-beam access imposes constraints on microscope and sample-stage geometry^28,29^, which impedes the use of a lift-out needle. Furthermore, these optics are incompatible with samples contained within a thick metal planchette, as is common for HPF. Recent work has introduced interferometric guidance without fluorescent fiducials or axial registration^30^, while a recent preprint combines depth correction with milled fiducials or fluorescence monitoring^29,31^. For HPF material, a central practical challenge is to maintain a local reference to rare targets through successive milling and handling steps on conventional integrated systems. Consequently, there remains a need for approaches that enable precise targeting of rare structures in thick specimens without requiring specialized microscope geometries, or extensive and potentially difficult-to-replicate ad-hoc modification of existing instrumentation. Here, we introduce two complementary approaches for fluorescence-guided targeting during cryo-FIB milling of thick cryogenic samples: with and without a lift-out needle. Both are implemented on a standard commercial cryo-FIB/SEM equipped with an integrated fluorescence light microscope (iFLM), without hardware modification or added fluorescent fiducial beads, and with no software beyond a Fiji^32^ script. These are: (i) iterative fluorescence correlation, in which repeated rounds of fluorescence imaging and milling progressively refine the target position; and (ii) bidirectional fluorescence correlation, which exploits the rotation of the sample during SOLIST to localize the target by fluorescence imaging from two orthogonal directions. We demonstrate the approaches using centrosomes in waffle-frozen early *C. elegans* embryos, a small and sparse target that represents the challenge of retaining a rare structure within a lamella of only a few hundred nanometres thick. As a benchmark, we compared it against an optimised SOLIST workflow, as this approach currently delivers the highest throughput of generated lamellae from HPF samples.

## RESULTS

### Centrosomes serve as a model for targeting rare subcellular structures

We evaluated fluorescence-guided targeting using centrosomes in early *C. elegans* embryos that at the one-to eight-cell stage contain only two to sixteen centrosomes. Using estimated volumes of approximately 4.2 µm³ per centrosome^33^ and 60,000 µm³ per embryo^34^, a single centrosome occupies approximately 0.007% of the total embryo volume. Retaining these small, sparsely distributed structures during lamella preparation therefore provides a test of subcellular targeting in high-pressure-frozen material.

To localize the centrosomes within isolated embryos, we used a strain expressing two fluorescent markers: GFP-tagged SPD-5, the primary scaffolding component of mature centrosomes^35,36^ and mScarlet-tagged NPP-10, which marked nuclear pores, allowing for visualisation of the nuclear envelope as a reference. Gravid adult *C. elegans* were bleached to release intact embryos (Fig. 1a), which were immediately waffle-frozen on EM grids and screened by cryo-light microscopy (Fig. 1b). We selected grids containing early embryos positioned away from grid bars, then processed them using iterative fluorescence correlation, bidirectional fluorescence correlation or a throughput-oriented, 2D-guided SOLIST workflow (Fig. 1c). A target-positive lamella was defined by recognizable centrosomes in TEM.

**Figure 1.**
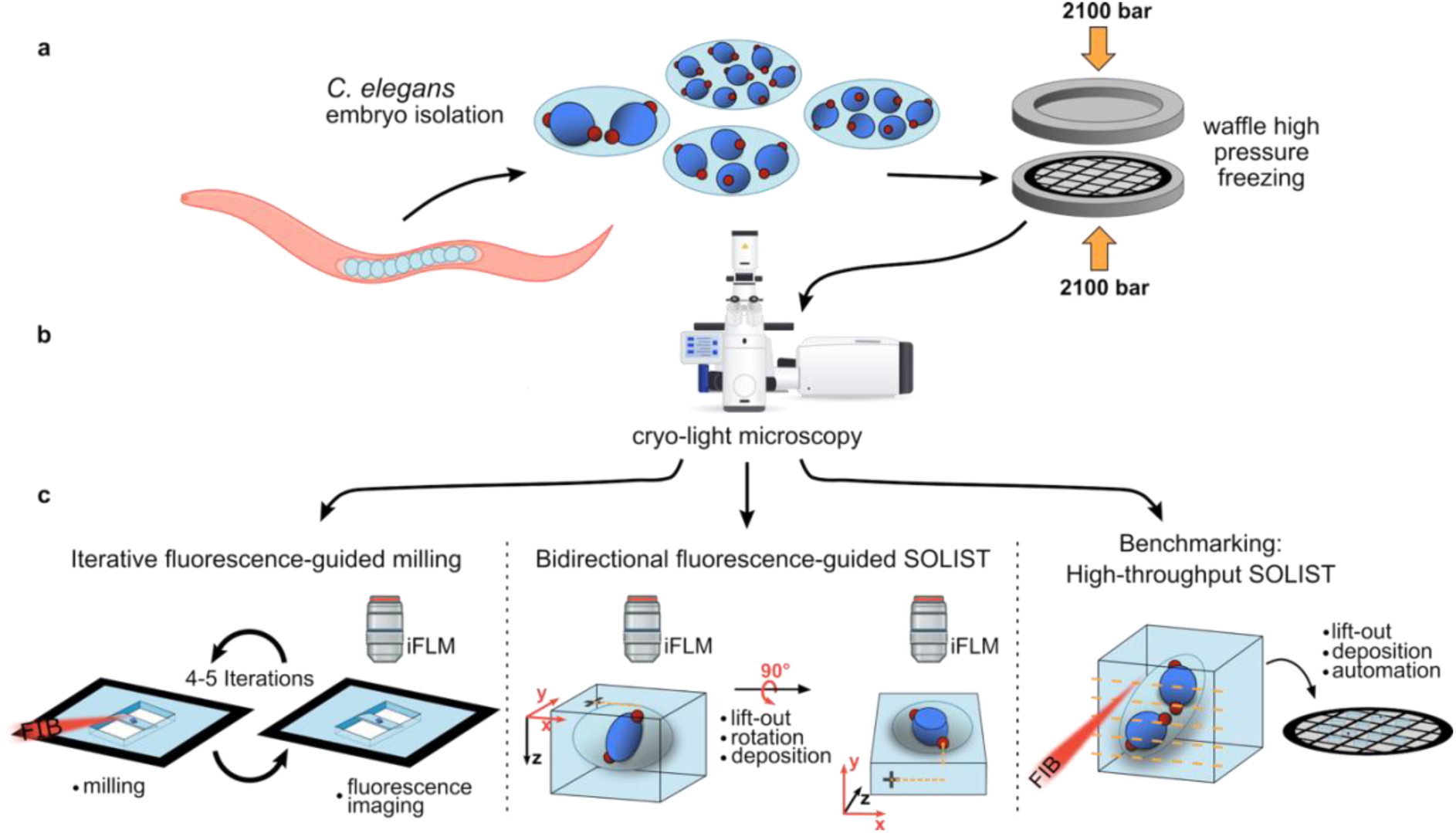
Three fluorescence-guided workflows for lamella preparation from waffle-frozen embryos. **a**) Isolation of *Caenorhabditis elegans* embryos followed by high-pressure freezing on electron microscopy grids using the waffle method. **b**) Screening of frozen grids by cryo-light microscopy to identify embryos and select target regions. **c**) Principles of the three workflows. Left, iterative fluorescence correlation alternates fluorescence imaging and focused ion beam (FIB) milling to refine target depth while thinning a slab directly on the original grid, without lift-out. Middle, bidirectional fluorescence correlation uses fluorescence images acquired before and after lift-out and deposition, which rotate the specimen by 90° relative to the iFLM optical axis, to locate the target from orthogonal directions. Right, 2D-guided high-throughput SOLIST produces serial slabs from a lift-out block; fluorescence-positive slices are selected for automated thinning using lateral target coordinates. Red spheres represent centrosomes and blue structures represent nuclei. iFLM - integrated fluorescence light microscopy.

### XY Correlation of the Cryo-Light and Electron Microscopy Images

All workflows began with correlation of fluorescence and SEM images to establish target position in the XY plane. Correlation began during sample preparation, when a pair of cooled fine tweezers was used to pierce a small hole through the ice in a single grid-square near the grid edge, providing a coarse registration landmark for correlating bright-field and fluorescence grid images with the SEM overview (Fig. 2a-d). We then refined the correlation locally at higher magnification using ice contamination on the grid bars as fiducial markers visible in SEM, FIB, and brightfield reflection images (Fig. 2e-h). For SOLIST workflows, holes in the support foil of the receiver grid provided additional local fiducial markers (Fig. 2i-l). To retain the target within the final lamella, we determined its depth within the frozen sample using either iterative fluorescence correlation or bidirectional fluorescence correlation.

**Figure 2.**
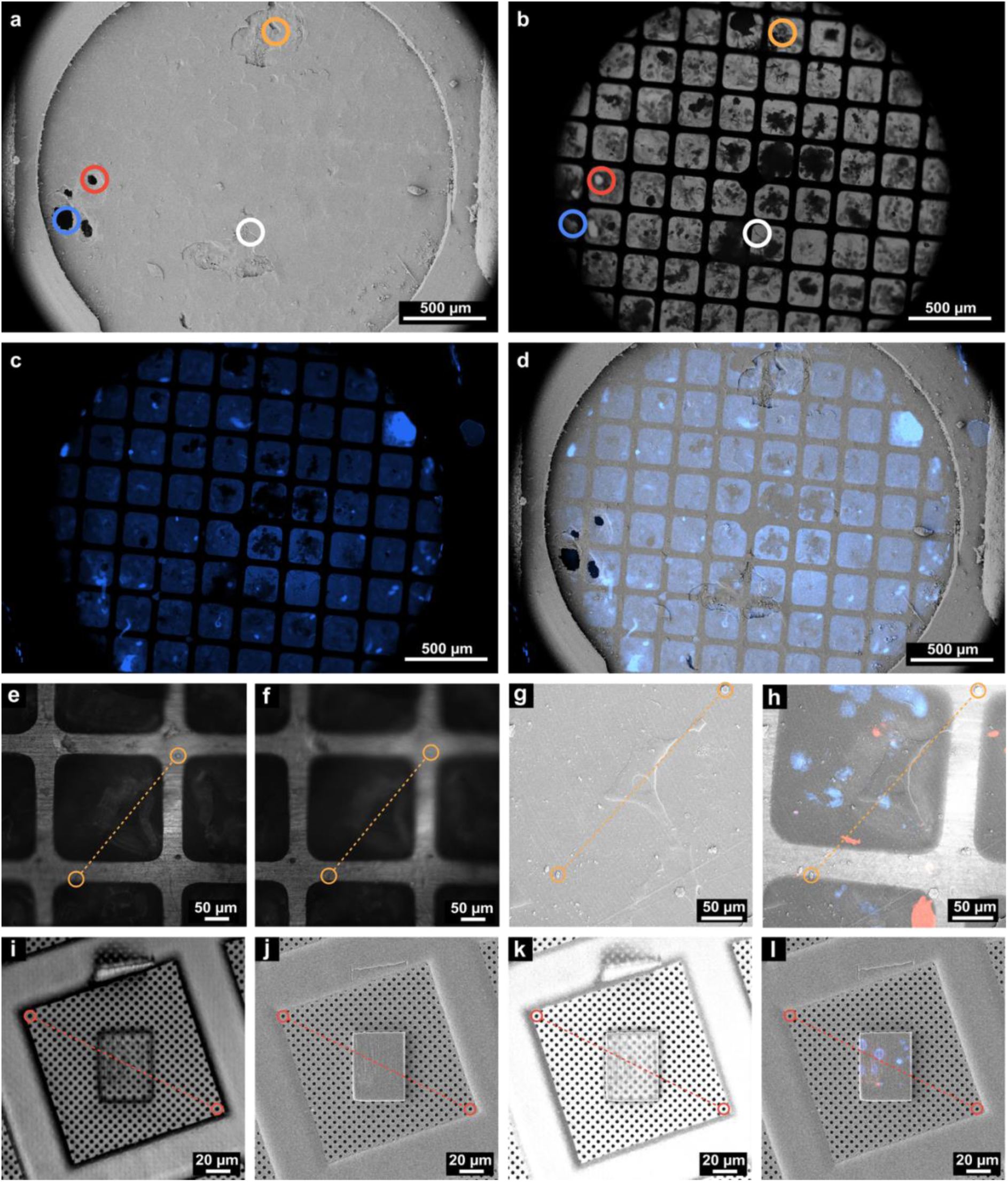
Correlation of fluorescence and SEM images from whole grids to individual deposited slices. **a-d**) Whole-grid correlation. **a**) Scanning electron microscopy (SEM) overview of a waffle-frozen grid. Coloured circles identify a hole pierced through the ice with cooled tweezers (blue), an air bubble (red), and distinctive surface features (white and orange). **b**) Transmitted-light overview acquired with a 5× objective, showing the corresponding landmarks. **c**) Fluorescence overview of the same grid. **d**) Fluorescence image registered to the SEM overview using the landmarks in a and b. **e–h**) Local correlation within a grid square. **e,f**) Reflection images acquired by integrated fluorescence light microscopy (iFLM) at two focal planes. Orange circles mark ice contamination on the grid bars used as local fiducials. **g**) SEM image of the same grid square, with the corresponding fiducials marked. **h**) Fluorescence channels overlaid on the registered SEM image. **i-l**) Correlation of a deposited SOLIST sliceon a receiver grid. **i**) iFLM reflection image; red circles mark holes in the support foil used as fiducials. **j**) SEM image of the same slicewith the corresponding holes marked. **k**) Registered overlay of i and j. **l**) Fluorescence channels overlaid on the SEM image. Fluorescence pseudocolours: SPD-5::GFP, red; NPP-10::mScarlet, blue. Scale bars, 500 µm (a-d), 50 µm (e-h) and 20 µm (i-l).

### Iterative Fluorescence Correlation - Targeting Without a Lift-Out Needle

To target centrosomes within the waffle-frozen embryos without a lift-out needle and with no equipment modification to a standard cryo-FIB-SEM system, we established a workflow relying on repeated fluorescence measurements during sequential milling.

We FIB-milled 50 x 50 µm trenches through the waffle-grid above and below the coordinates of the ROI determined by the initial XY correlation step. Milling trenches is part of standard waffle-grid preparation^22^ as it enables access to the target region both for thin lamella preparation during FIB-milling and for tilt series acquisition during TEM imaging. After milling the undercut, the resulting volume of biological material was approximately 15-µm thick, hereafter referred to as the slab to distinguish it from the final polished lamella (Fig. 3a). Using an integrated fluorescence light microscope (iFLM), we acquired a wide-field fluorescence z-stack of the slab using an axial step size of 0.5 µm or 0.35 µm^37^ (Fig. 3b). We computationally resliced the z-stack from XY into YZ planes and rotated by the milling angle to visualise the fluorescence signal relative to the slab surfaces (Fig. 3c). Within the re-sliced stacks, we estimated the position of the upper front edge of the slab using the strong reflection signal provided by the GIS layer deposited on the front surface (Fig. 3d). Assuming a cuboidal slab, we extrapolated the top surface across the slab at this z height (Fig. 3e). We defined the bottom surface of the slab as the plane parallel to the top surface, offset by the slab thickness either measured from FIB imaging or estimated from the user-specified milling parameters (Fig. 3f).

**Figure 3.**
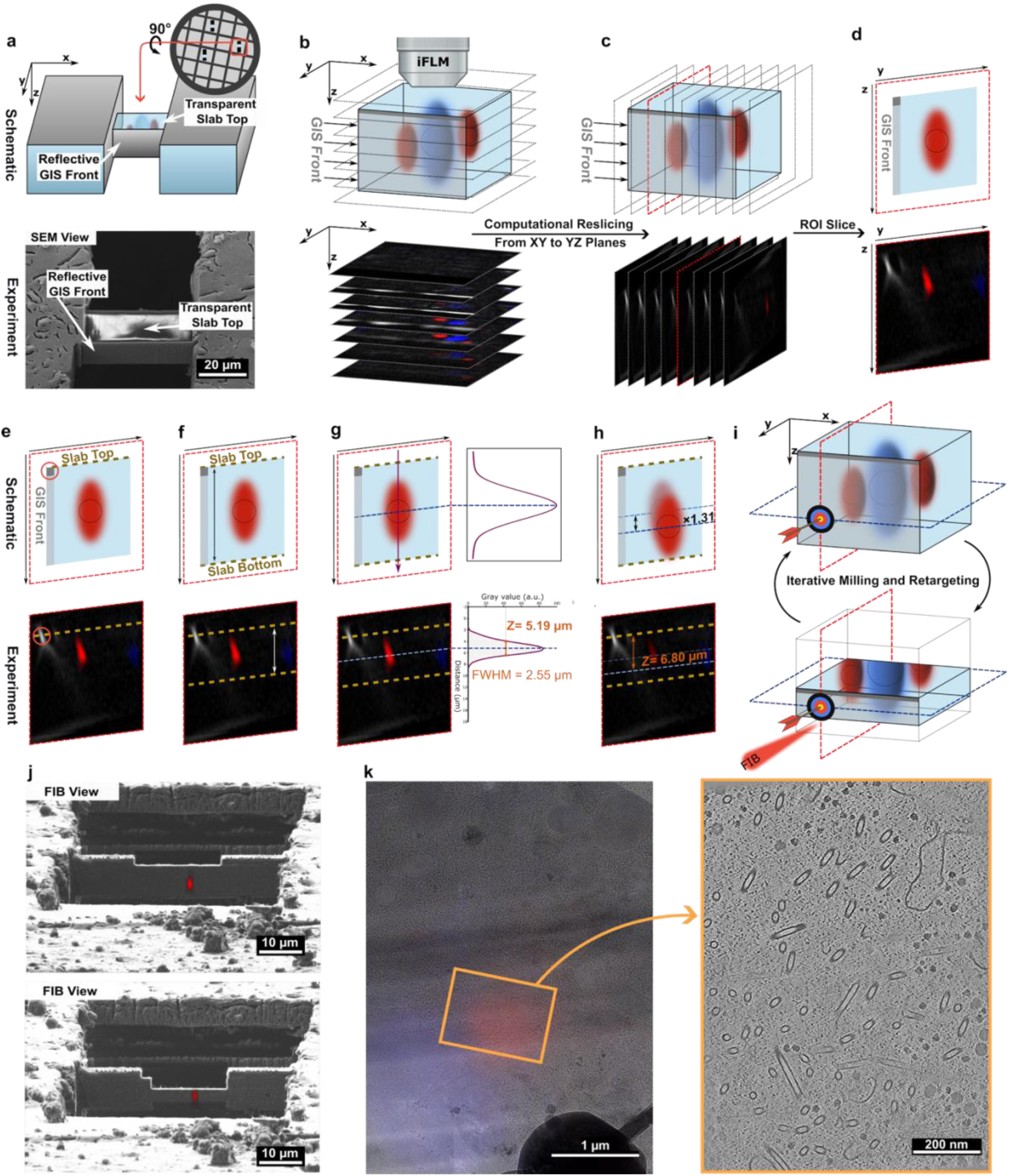
Iterative fluorescence correlation guides target retention during on-grid lamella thinning. **a-i**) Workflow illustrated by schematics and corresponding experimental images. **a**) An undercut slab on a waffle-frozen grid, with a reflective protective layer on its front face and an exposed upper surface for fluorescence imaging. **b**) Acquisition of a fluorescence z-stack using integrated fluorescence light microscopy (iFLM). **c**) Computational reslicing of the XY stack into YZ planes, followed by rotation by the milling angle to align the slab surfaces. **d**) Selection of the resliced plane containing the region of interest (ROI). In the schematics, the elongated fluorescence distribution illustrates axial broadening, whereas the outlined structure indicates the underlying target. **e**) Identification of the upper front edge of the slab from the reflection signal of the protective layer deposited by the gas injection system (GIS; circle). This reference defines the upper surface of the slab. **f**) Definition of the lower surface using the slab thickness measured by focused ion beam (FIB) imaging or specified in the milling parameters. **g**) Estimation of the fluorescence centre from the midpoint between the two half-maximum positions of the axial intensity profile. The blue dashed line marks this centre; FWHM, full width at half maximum. **h**) Multiplication of the apparent target depth by 1.31 to approximate the correction for refractive-index mismatch between vacuum and vitreous ice. **i**) Repeated depth measurement and milling to progressively thin the slab around the target. **j**) Fluorescence-FIB overlays at two successive stages of thinning. **k**) Fluorescence overlaid on a transmission electron microscopy (TEM) lamella map (left); the orange box marks the region shown as a tomographic slice (right). Fluorescence pseudocolours: SPD-5::GFP, red; NPP-10::mScarlet, blue. Scale bars, 20 µm (a), 10 µm (j), 1 µm (k, left) and 200 nm (k, right).

The anisotropic point spread function of epifluorescence microscopes results in significantly poorer axial than lateral resolution and produces a signal elongated along the Z axis. However, for an isolated fluorescent signal, the intensity profile along the Z axis can be approximated by a Gaussian-like distribution, allowing the center of the signal to be estimated despite its substantial axial broadening. We estimated the Z position of the fluorescent target as the midpoint of the full width at half maximum (FWHM) of the intensity profile (Fig. 3g). Finally, we estimated the target depth within the imaged slab as the apparent distance between the FWHM midpoint and the upper surface of the slab multiplied by 1.31, as an approximate correction for the refractive-index mismatch between vacuum and vitreous ice^19^ (Fig. 3h). Although this correction factor depends in principle on the angle between the optical axis and the sample surface, which here equals the milling angle, the effect is small: at a milling angle of 10°, the factor differs from that for a perpendicular surface by less than 1%^38^. We used this depth estimate to guide iterative rounds of FIB milling and imaging (Fig. 3i). We performed image reslicing and depth calculation using a Fiji^32^ script that includes an optional tool to overlay the fluorescence signal onto the FIB image at the milling angle to facilitate targeting (Fig. 3j).

After material was removed from either side of the target, a new stack provided an updated position relative to the remaining slab (Fig. 3i,j). Re-measuring this local reference allowed us to adjust for residual localization errors, stage drift and ion beam-induced sample distortion, which can make single-step 3D-correlation too imprecise to retain the target in the final lamella^27^. Thus, the established workflow relies on iterative rounds of imaging, correlating and milling rather than completing the targeting in a single pass. We gradually reduced the amount of material removed at each iteration as the milling patterns approached the target. Approximately five iterations were usually sufficient to progress from a 15-µm slab to a thickness of 1 µm. The additional imaging and correlation required approximately 5–10 min per iteration. Below a sample thickness of 1 µm, the signal is usually too weak to produce an intensity profile from which a Gaussian-like peak can be seen, hence the target centre cannot be reliably determined. We therefore did not iterate beyond this point, and instead polished to the final lamella thickness using the target position determined in the last iteration. Additionally, we introduced micro-expansion joints at approximately 3 µm thickness to limit slab bending during subsequent thinning^39^. We placed these stress relief cuts near the estimated final height of the lamella. Delaying the cuts until a slab thickness of approximately 3 µm allowed further refinement of the target position, whereas further thinning without the cuts could cause excessive bending and breakage (Supplementary Fig. 1). These steps produced lamellae with a nominal final thickness of 200 nm.

We prepared six lamellae across two grids, with three sites per grid. All six survived transfer to the TEM, and five contained recognizable centrosomes (Fig. 3k and Supplementary Fig. 2a; Table 1). Preparing a grid with three polished lamellae from waffle-frozen samples required approximately 39 hours of operator time, or 13 hours per lamella with a nominal thickness of 200 nm. However, most of this time comes not from the iterative milling steps, but from milling the large initial trenches, performing the undercut, and the XY correlations described above. These time-intensive steps are common for any milling workflow aimed at producing thin lamellae directly on waffle frozen samples, whether or not iterative targeting is used. Preparation from the approximately 15 µm thick slab to the final polished lamella took an average of 4 hours, including an average of five targeting iterations per site.

**Table 1.** Preparation times and target recovery rates for iterative fluorescence correlation, bidirectional fluorescence correlation, and 2D-guided high-throughput SOLIST.

|  | **Iterative fluorescence correlation** | **Bidirectional fluorescence correlation** | **2D-guided high-throughput SOLIST** |
|---|---|---|---|
| **Milled sites** | 6 | 9 | 20 |
| **Polished lamellae** | 6 | 9 | 10 |
| **Transfer survival to TEM** | 6/6 (100%) | 6/9 (67%) | 8/10 (80%) |
| **Target-positive TEM-imaged lamellae** | 5/6 (83%) | 4/6 (67%) | 2/8 (25%) |
| **Overall target yield per attempted preparation** | 5/6 (83%) | 4/9 (44%) | 2/10 (20%) |
| **Total time** | 78 hours | 54 hours | 36 hours |
| **Operator time** | 78 hours | 54 hours | 26 hours |
| **Automated time** | 0 hours | 0 hours | 10 hours |
| **Average time from slab/slice to lamella** | 4 hours | 2 hours | 1 hour |
| **Average time from slab/slice to hit target** | 4.8 hours | 4.5 hours | 5 hours |
| **Average time per lamella** | 13 hours | 6 hours | 3.6 hours |
| **Average time per hit target** | 15.6 hours | 13.5 hours | 18 hours |

### Bidirectional Fluorescence Correlation - Targeting with a Lift-Out Needle

For FIB-SEM systems equipped with a lift-out needle, we developed a complementary targeting approach based on the SOLIST workflow^8^. We combined fluorescence images of the ROI acquired from orthogonal directions before and after slice deposition to determine its position and guide FIB milling. Unlike conventional SOLIST, which aims at generating multiple lamellae from each lift-out block, our bidirectional targeting workflow uses each block to prepare a single lamella targeted to the selected ROI.

After the initial fluorescence-SEM correlation (Fig. 4a), we FIB-milled a cross-shaped marker at low ion-beam current approximately 10 µm from the target along the X axis, to mark its position in the XY plane (Fig. 4b). We then positioned the milling patterns around the target for lateral trench milling to enable lift-out of an 18 × 30 µm block (Fig. 4c-d).

**Figure 4.**
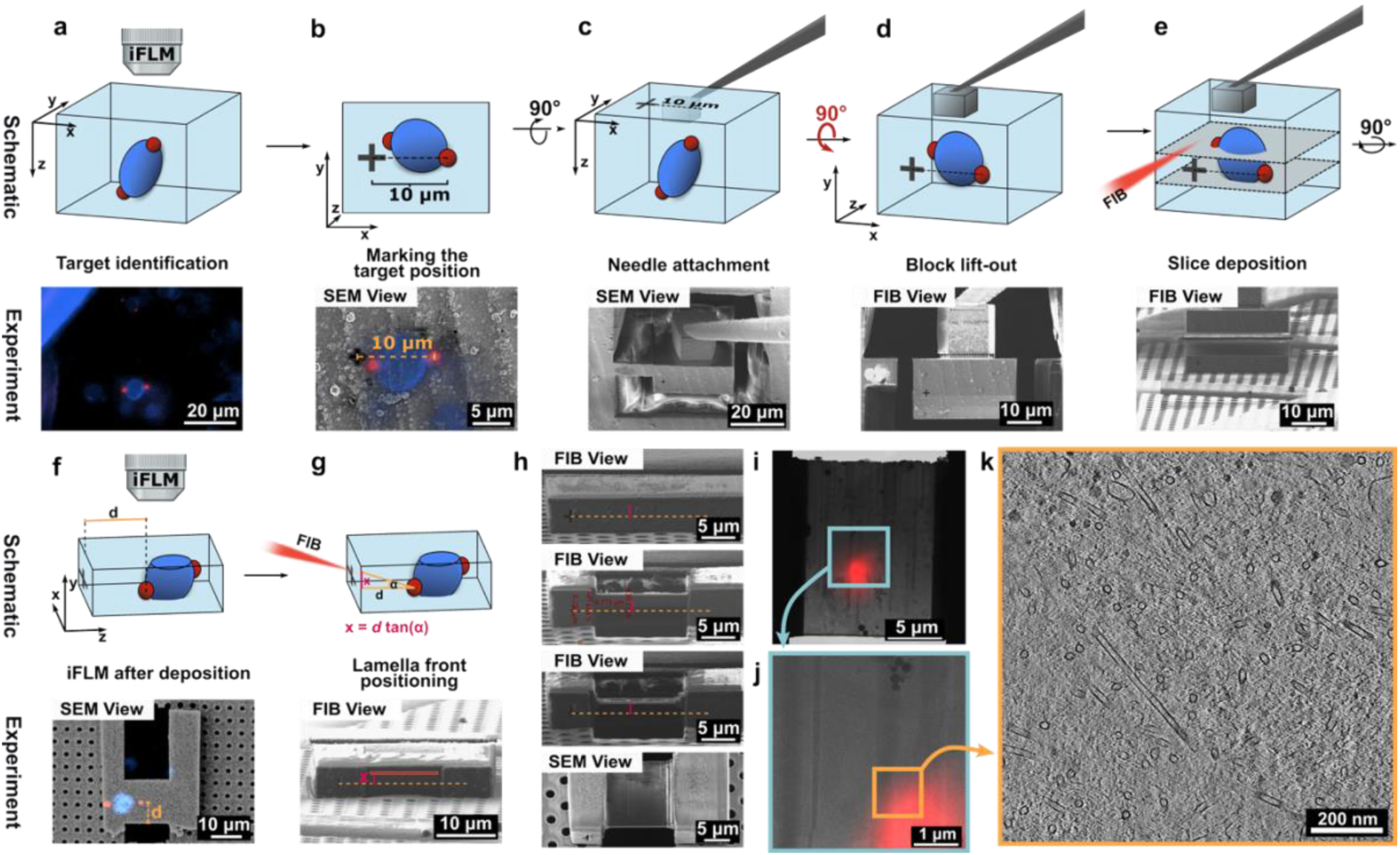
Bidirectional fluorescence correlation uses orthogonal views to position a lamella around the target. **a-g**) Workflow illustrated by schematics and corresponding experimental images. **a**) Initial identification of the fluorescent target in a waffle-frozen embryo. **b**) Correlation of fluorescence and scanning electron microscopy (SEM) images, followed by milling of a cross-shaped reference marker approximately 10 µm from the target along the X axis. **c**) Milling around the selected block and attachment of the lift-out needle. **d**) Block release and lift-out; the cross remains visible in the focused ion beam (FIB) image as a reference for target height. **e**) Deposition of a slab containing the target onto a receiver grid. **f**) Fluorescence imaging after deposition provides an orthogonal view because the specimen has rotated by 90° relative to the iFLM optical axis. The distance d is measured from the cleaned slab front to the target centre. **g**) Positioning of the lamella front using the offset x = d tan(α), where α is the milling angle and x is the displacement from the target height indicated by the cross. **h**) Successive stages of milling guided by the cross and calculated offset. The upper three panels are FIB images; the lower panel is an SEM image of the polished lamella. **i**) Transmission electron microscopy (TEM) lamella map overlaid with the fluorescence image acquired after slab deposition. **j**) Enlargement of the turquoise-boxed region in i; the orange box marks the region shown as a tomographic slice in **k**. Fluorescence pseudocolours: SPD-5::GFP, red; NPP-10::mScarlet, blue. iFLM, integrated fluorescence light microscopy. Scale bars, 20 µm (a,c), 5 µm (b,h,i), 10 µm (d–g), 1 µm (j) and 200 nm (k).

On the lifted block, the cross-shaped marker is visible by FIB imaging and indicates the height of the ROI within the block (Fig. 4d). We used this axial reference to guide the deposition of an approximately 4 µm thick slice containing the selected region onto the receiver grid (Fig. 4e), thus reducing the risk of unintentionally milling away the target during this step. Following deposition, we re-imaged the slices using iFLM. Transfer to the receiver grid rotated the block by 90° relative to the iFLM optical axis (Fig. 4a,f). Re-imaging therefore provided an orthogonal view of the same target: the dimension initially aligned with the optical Z axis was now imaged in the iFLM XY plane, allowing that coordinate to be determined using the higher lateral resolution of the microscope. We correlated the newly acquired fluorescence stack with the SEM image of the deposited slices to establish the position of the target within them (Fig. 4f). Using FIB-milling, we cleaned the front surfaces of the deposited slices and removed the excess material posterior to the target to reduce the distance over which angular errors could accumulate (Supplementary Fig. 3a-d). We then measured the distance *d* from the cleaned front of the slice to the center of the fluorescence signal in the slice XZ plane (Fig. 4f). Because the final lamella is inclined relative to this plane, its front must be offset from the cross-defined target height to intersect the target further back in the slice. We calculated this offset as (Fig. 4g):

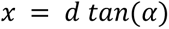

where *α* is the user-defined milling angle in degrees, *d* is the distance from the front of the slice to the center of the target in the XZ plane, and *x* is the vertical offset of the lamella front from the reference height indicated by the cross-shaped marker (Fig. 4g). Having determined *x*, we positioned the lamella front accordingly to perform rough milling. The cross served as a persistent indicator of the target height, allowing the milling patterns to be checked throughout the manual milling process and their positions to be adjusted when necessary to compensate for stage drift or misalignment between different ion-beam currents (Fig. 4h).

The geometric correction assumes that the slice lies flat on the support foil. Uneven foil or contamination beneath a deposited slice changes its orientation relative to the ion beam and can displace the predicted lamella position (Supplementary Fig. 3e). We therefore deposited slices on clean, flat foil.

We prepared nine lamellae using bidirectional fluorescence correlation. Six survived transfer to the TEM, and four contained recognizable centrosomes (Fig. 4i–k and Supplementary Fig. 2b; Table 1). Carrying out the bidirectional fluorescence correlation workflow required a total of 54 hours, corresponding to an average time of 6 hours per lamella and 13.5 hours per recovered target. Following slice deposition, iFLM imaging, GIS coating, front and back cleaning, correlation, manual milling and polishing together took approximately 2 hours per finished lamella.

### Benchmarking Against 2D-Guided High-Throughput SOLIST

To compare the two targeting workflows with current state-of-the-art lift-out practices, we used a SOLIST^8^ implementation that prioritized serial slice production and automated lamella thinning while retaining fluorescence guidance in XY (Fig. 5a-b). This comparator selected fluorescence-positive slices and used their lateral fluorescence coordinates to guide preparation, without applying the iterative depth measurements or geometric targeting methods described above. We also refined the preparation procedure to improve lamella quality and mechanical stability. Slices were deposited on non-sputter-coated UltrAuFoil grids to improve their attachment to the foil, thus greatly reducing the lamella loss rate. Using uncoated receiver grids, the micro-stitches aimed at stabilizing the slices, as proposed in the original SOLIST paper^8^, were not required, as we observed no improvement in attachment when they were included (Supplementary Fig. 4). Additionally, to deposit the slices on the receiver grid more robustly, we optimized the deposition patterns. We released slices from the block using a thin cleaning cross-section pattern, scanning from left to right of the block, immediately followed by a line pattern. This method limits the reattachment of the slice to the block caused by redeposition and promotes its electrostatic deposition on the foil of the receiver grid. With these refinements, the slice deposition rate increased, with each slice deposition taking approximately 3-5 minutes.

**Figure 5.**
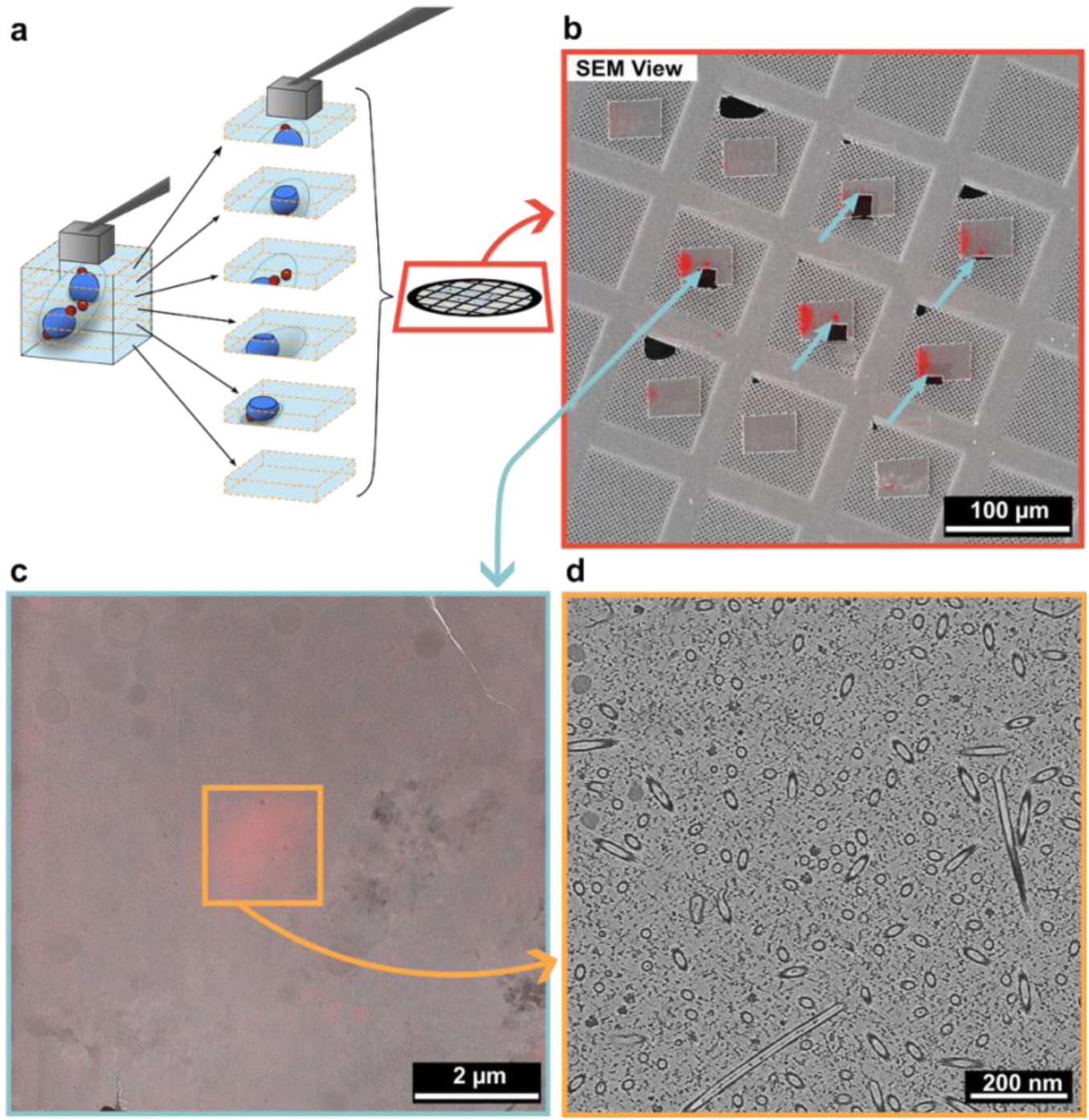
2D-guided high-throughput SOLIST combines serial slab production with fluorescence-based selection and lateral targeting. **a**) Schematic of serial slab deposition from a lift-out block. Only a subset of slabs contains the target, represented by red spheres; blue structures represent nuclei. **b**) Fluorescence overlaid on a scanning electron microscopy (SEM) overview of deposited slabs. Turquoise arrows indicate fluorescent targets in selected slabs. The slab fronts have been trimmed to reduce the distance between the target and the milling front. **c**) Fluorescence image of a polished lamella overlaid on its transmission electron microscopy (TEM) map. **d**) Tomographic slice from the orange-boxed region in c, showing a centrosome. Fluorescence pseudocolours: SPD-5::GFP, red; NPP-10::mScarlet, blue. Scale bars, 100 µm (b), 2 µm (c) and 200 nm (d).

We also observed that the top surface of deposited slices was often uneven. Consequently, application of a GIS layer from a single, normal-incidence direction could produce a non-uniform protective layer and an irregular milling front, increasing lamella breakage (Supplementary Fig. 5a-b). We therefore applied GIS in two shorter steps, one at the mapping position and one with the sample held near perpendicular to the ion beam (See Methods). This approach allowed us to coat shadowed regions, producing a more homogeneous GIS protective layer (Supplementary Fig. 5c). Each slice was then reduced in size by removing material from the leading and trailing ends of the slice with the sample held perpendicular to the ion beam, so that the new front edge lay closer to the ROI. We observed that smaller lamellae were generally more stable and less prone to breakage. A clean slice front also provides a smoother surface for the final GIS deposition, protecting the slice during subsequent milling. In addition, when applying the protective GIS layer on the slice front, we rotated the stage so that the GIS needle pointed perpendicular to the slice front, which resulted in increased thickness and uniformity of the protective layer at the leading edge of the lamella. Finally, we thinned the deposited slices by automated milling with AutoTEM Cryo (Thermo Fisher Scientific) overnight to maximize effective instrument time.

For the comparison with the targeted approaches, we applied this refined workflow to the same *C. elegans* embryo samples. After the initial fluorescence-SEM correlation, we selected embryos with 4-8 centrosomes and adjusted the size of the lift-out block to cover the entire embryo. Without further targeting, we sliced the lifted block into slices of even thickness (commonly 10 slices per block). We deposited twenty slices from two embryos at approximately 5 µm intervals. After deposition, we acquired iFLM images of the slices that we overlaid and registered with the SEM images to confirm the presence of fluorescent targets and to determine their approximate positions (Fig. 5a-b). For this step, we used only the two-dimensional target position in the slice XY plane to guide subsequent lamella preparation; no three-dimensional correlation was performed. Ten of the twenty slices contained fluorescent ROIs, and were selected for automated overnight milling. Eight of the ten lamellae survived transfer into the transmission electron microscope, and two of these contained centrosomes, resulting in a target-positive fraction of 25% among the surviving lamellae (Table 1, Fig. 5c-d, Supplementary Fig. 2c).

The 2D-guided high-throughput SOLIST workflow required a total of 36 h, comprising 26 hours of operator time and 10 hours of automated milling (Table 1). This corresponded to an average of 3.6 hours per polished lamella and 18 hours per recovered target. Preparation from a deposited slice to a polished lamella took an average of 1 hour.

### Target yield and instrument use across workflows

To compare the three approaches, we recorded the proportion of lamellae containing centrosomes and the total preparation time required to prepare them (Fig. 6a-c). Among lamellae imaged in TEM, the target-positive fractions were 83% for iterative fluorescence correlation (5/6 lamellae), 67% for bidirectional fluorescence correlation (4/6 lamellae) and 25% for 2D-guided high-throughput SOLIST (2/8 lamellae) (Fig. 6a). Microscope time per lamella was 13 hours, 6 hours, and 3 hours 36 minutes, respectively (Fig. 6b). Per recovered target, the corresponding times were 15 hours 36 minutes, 13.5 hours, and 18 hours (Fig. 6c). Recent developments, such as plasma Focused Ion Beams (pFIBs)^40^ and non-devitrifying femtosecond ablative laser pulses^41^ may reduce the time taken in bulk milling steps. We therefore also compared the time required to prepare a final polished lamella, starting from a deposited slice for the lift-out methods or from an undercut slab for iterative fluorescence correlation. Using this metric, the milling times per polished site were 4 hours for iterative correlation, 2 hours for bidirectional correlation, and 1 hour for 2D-guided high-throughput SOLIST (Fig. 6d). Dividing the total time spent on these steps by the number of recovered targets gave 4 hours 48 minutes for the iterative approach, 4 hours 30 minutes for bidirectional correlation, and 5 hours for high-throughput SOLIST (Fig. 6e).

**Figure 6.**
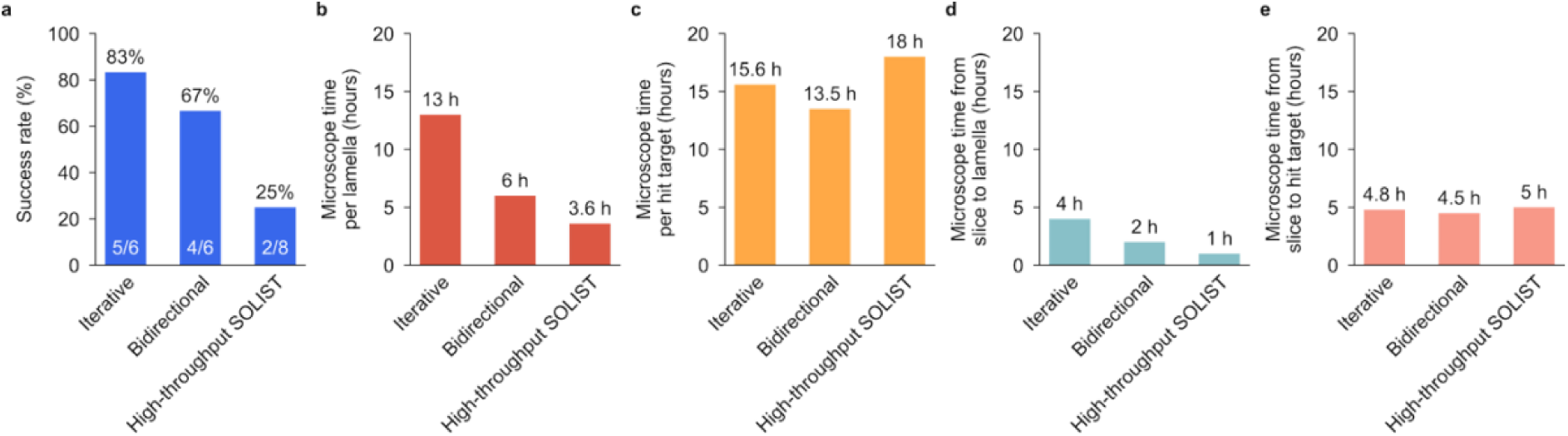
Target recovery and preparation time across the three workflows. In each panel, workflows are ordered as iterative fluorescence correlation, bidirectional fluorescence correlation and 2D-guided high-throughput SOLIST. **a**) Fractions of lamellae examined by transmission electron microscopy (TEM) that contained recognizable centrosomes: 5/6 (83%), 4/6 (67%) and 2/8 (25%), respectively. Fractions within the bars give the number of target-positive lamellae divided by the number examined in TEM; lamellae lost before TEM examination are excluded from this denominator. **b,c**) Total preparation time divided by the number of polished lamellae (**b**; n = 6, 9 and 10) or target-positive lamellae recovered in TEM (**c**; n = 5, 4 and 2). Total preparation times were 78, 54 and 36 h, respectively, and include automated milling where used. **d**) Mean preparation time from the undercut slab for iterative fluorescence correlation, or from a deposited slice for the two SOLIST workflows, to a polished lamella. **e**) Total time spent on the preparation steps represented in d divided by the number of target-positive lamellae recovered in TEM. A recovered target denotes a target-positive lamella. Bars show descriptive values for the preparations summarized in Table 1.

## DISCUSSION

Keeping a rare target within a lamella requires its position to be tracked as the surrounding specimen is progressively removed. Established methods for three-dimensional cryo-CLEM guided milling rely on bead-based correlation for retaining fluorescent ROIs in the final lamella^26^ or use coincident microscopes that allow fluorescence imaging during ion-beam milling without stage movement^28,29^. Recent interferometric guidance enables target preservation without fluorescent fiducials or axial registration^30^ and a recent preprint^31^ also combines correction for refractive-index effects with milled fiducials or fluorescence monitoring for targets of different sizes. Although control software for FIB-SEM microscopes such as fibsemOS can now correct refractive-index mismatch, bead-based workflows that rely on a single initial correlation remain susceptible to axial drift and localization or registration errors. Therefore, tri-coincident systems provide the most direct feedback on target position by on-the-fly imaging, however, they currently require extensive instrument modification^28,29^, are not commercially standardized, and impose geometric constraints that can prevent lift-out. Moreover, confirming that the target has been reached generally depends on observing changes in its fluorescence during milling, which entails removing part of the ROI.

To overcome the limitations of current workflows, we developed two complementary methods to target rare ROI within high-pressure-frozen material using commercial cryo-FIB-SEM equipped with an integrated fluorescence microscope. This provides additional options for preparing high-pressure-frozen specimens without introducing fluorescent beads or requiring simultaneous access for fluorescence imaging and ion-beam milling. The iterative fluorescence correlation workflow follows target depth through repeated measurements relative to the remaining slab, while the bidirectional fluorescence correlation method instead uses orthogonal optical views and a persistent milled reference to position the lamella. Both approaches positioned the lamella before the target is reached and recovered centrosomes from waffle-frozen *C. elegans* embryos, providing reliable routes to targeted preparation with and without a lift-out needle.

Apart from a simple Fiji script, no additional software is required, so the approaches can be adopted by a broad range of laboratories without specialized training or costly installations. Just as importantly, they are robust, as two features make these approaches tolerant of the errors that affect single-pass correlation. First, the reference point lies close to the ROI: either the GIS-coated top of the slice or a cross milled a few micrometres from the target. Because angular and scaling errors grow with distance from the reference, this limits their effect on the final lamella position. Second, the reference remains available throughout milling. In iterative fluorescence correlation the target is re-localized at every round, and in bidirectional fluorescence correlation the cross can be checked at each milling step, so drift, slice deformation and misalignment between milling currents can be corrected before they compromise the lamella.

Aside from providing the highest target accuracy, the main advantage of the iterative fluorescence correlation workflow is that it offers a way to target specimens when a lift-out needle is unavailable, enabling robust targeted milling of high-pressure-frozen material for laboratories that have so far worked only with plunge-frozen samples. It is also applicable to advanced instruments where a lift-out needle cannot be installed, such as the Arctis, which has reported much faster rates of milling through HPF samples than a gallium FIB^40^. However, the reliability of the depth estimate depends on fluorescent signal strength, separation from neighbouring features and how closely the fluorescent label represents the structure of interest. With the highest target recovery rate among the tested workflows, this approach is well suited to rare targets, where each lamella site represents a substantial investment of time. However, the method cannot be applied to HPF samples vitrified within planchettes and its application is restricted to waffle-frozen samples, thus limiting sample thickness to approximately 25-35 µm. Repeated measurements allowed us to update target position if the slab moved or deformed during milling, provided that the label remained bright enough through successive imaging rounds. Although photobleaching is usually diminished at cryogenic temperatures^42,43^, it is not entirely eliminated and thus very sensitive tags may suffer. Extensive SEM imaging may exacerbate this problem^44^ and should thus be avoided. Finally, the iterative approach relies on isolated, discrete signals; as fluorescence overlaps from several structures render localization of a single target unreliable.

In contrast, the fiducial cross milled in bidirectional fluorescence correlation is insensitive to z drift, so multiple iterations of fluorescence imaging are unnecessary. Although the method’s accuracy still depends on the initial XY correlation, the orientation of the deposited slice and the milling angle, its success rate (67%) is comparable to that of iterative fluorescence correlation (83%), making it suitable for targeting rare targets. Beyond requiring a lift-out needle, the main limitation of this approach is its sensitivity to deviations between the nominal and actual milling angle. A slice resting on debris or on distorted foil will be tilted relative to the ion beam. An angular error of only 0.5°, difficult to see by eye alone, can then displace the lamella by up to ∼175 nm at the target, depending on the target’s distance from the block front. Since the target must remain within a ∼200 nm thick lamella, such an offset can be enough to lose it. Slices should therefore be deposited onto clean, levelled foil, and the block front trimmed to place the slice front closer to the target, thus limiting the distance over which an angular error can shift the intended lamella plane. Additionally, the method requires accurate initial 2D correlation between the fluorescence and SEM data, as this determines the placement of the reference mark. Adopting the method thus requires access to suitable lift-out hardware and expertise.

We benchmarked the two targeted approaches against a 2D-guided SOLIST workflow that incorporates several protocol refinements and, in our hands, provides the highest throughput for lamella preparation from high-pressure-frozen samples. Although high-throughput SOLIST prioritizes lamella production over precise targeting, our implementation was guided by fluorescence. Compared to the high-throughput SOLIST workflow, both targeted approaches retained the target considerably more often. However, the small datasets call for caution, as transfer losses greatly affect the overall yields per attempted preparation (Table 1). We scored success by the presence of recognizable centrosomes in TEM. This establishes recovery of the target region, but does not measure how much of an individual centrosome was retained or the distribution of three-dimensional localization errors. Precise depth targeting is likely to be most valuable when a specimen contains only a few sites of interest that can each be localized reliably. Serial preparation in such specimens can consume substantial effort on planes that never intersect the target. On the other hand, for abundant targets, or targets extending through larger volumes, fluorescence-based slice selection followed by automated thinning can provide sufficient recovery with less manual intervention. In conclusion, where the balance lies will depend on target size and distribution, labelling quality and specimen geometry.

Time is a key consideration when comparing approaches, as access to cryo-FIB instruments is often limited. Timing also needs to be interpreted in relation to the outcome being measured. While high-throughput SOLIST was the fastest route to polished lamellae, accounting for target recovery narrowed the differences in total preparation time, to 15.6, 13.5 and 18 h per recovered target for iterative correlation, bidirectional correlation and high-throughput SOLIST, respectively. From the exposed slab or deposited slice onward, the corresponding times were 4.8, 4.5 and 5 h per recovered target (Fig. 6). Both the starting geometry and the amount of material removed contribute to these values, alongside targeting overhead. For iterative correlation, bulk milling and the initial correlation accounted for much of the preparation time, with each subsequent imaging and correlation cycle taking approximately 5-10 min. Operator time is another consideration, as the 36-h high-throughput SOLIST workflow included 10 h of automated milling. Centrosomes are also a favourable case for high-throughput SOLIST, as each selected embryo contained four to eight; for structures rarer within a grid or lift-out block, its time per hit target would rise accordingly. Choosing a workflow therefore means weighing instrument occupancy and operator time against the number of targets recovered.

The changes we made to SOLIST may also be useful beyond the present comparison. Deposition on receiver grids without sputter coating, modified release patterns, GIS application from two directions and trimming of the slice ends supported serial preparation followed by automated thinning, increasing the throughput of the originally published method. Under the conditions tested, we could also omit micro-stitches and simplify preparation, although attachment behaviour will need to be reassessed for other grids and specimens. The use of thick silver coatings to strengthen attachment to support films in other SOLIST implementations^13^ illustrates how these choices depend on the receiving substrate. Lamella size should likewise reflect the biological question, balancing target retention with preservation of the surrounding context. When the target occupies only a small region, removing unnecessary material can simplify the preparation of a smaller lamella centred on it. A larger lamella may be preferable when the aim is to retain the surrounding cellular architecture or sample several sites on a single lamella.

The choice of preparation strategy also needs to account for how targets will be located and verified in the final lamella. In the high-throughput experiment, fluorescence remained detectable after polishing and guided TEM acquisition, allowing us to locate candidate regions without an exhaustive TEM search. For weakly labelled structures, or those without a recognizable ultrastructural signature, confirmation may require further on-lamella correlation^45^. The distinction between using fluorescence to position a lamella and establishing the identity of the retained structure applies to all three workflows.

Further gains in efficiency may come from reducing the bulk milling that precedes precise targeting. Plasma ion beams^40^ and the laser-ablation approach described in a recent preprint^41^ could accelerate material removal, although their integration with the present workflows remains to be tested. Automating detection of the GIS reference, assessment of fluorescence profiles and transfer of target coordinates into milling patterns could also reduce manual intervention. Estimates of positional uncertainty would help determine when a further measurement is needed before more material is removed. Extending these workflows beyond the centrosome model to tissues, organoids and other fluorescent targets will require testing across differences in optical background, labelling density and mechanical behaviour.

Our results provide complementary ways to retain selected centrosomal regions during lamella preparation from high-pressure-frozen embryos. The most suitable workflow depends on the experimental question and the balance between targeting precision and throughput. Hardware availability, sample type, target labelling, abundance and cellular location also influence this choice. Among the workflows tested, iterative fluorescence correlation alone enables targeted milling of waffle-frozen material without a lift-out needle. It does, however, require bright, discrete and photostable labelling, as well as substantial microscope time. Where lift-out hardware and expertise are available, bidirectional fluorescence correlation reaches the target with fewer imaging rounds and less milling, although its accuracy remains sensitive to deposition geometry. Both approaches are particularly valuable for rare targets. When targets are abundant, or the aim is to acquire large datasets without selecting specific structures, the additional time and effort required for precise targeting may offer limited benefit. Throughput-oriented workflows such as SOLIST may then be the more efficient choice.

Selecting or combining these workflows allows preparation to be tailored to target rarity, sample value and the available instrumentation. As in situ structural biology expands to more complex specimens, including high-pressure-frozen tissues and organoids, recovering rare structures within their native cellular context is likely to become increasingly important. The approaches presented here can be implemented on existing commercial cryo-FIB-SEM systems and provide a practical foundation for extending targeted cryo-ET to a broader range of fluorescently labelled structures and biological specimens.

## METHODS

### C. elegans culture, staging and embryo preparation

Adult *C. elegans* worms from a line expressing SPD5::GFP and NPP10::mScarlet were cultured at 20 °C on nematode growth medium (NGM) plates seeded with *Escherichia coli* OP50, as described^46^. For HPF preparation synchronized young adults were obtained by a timed egg lay. Ten 6 cm NGM plates were each seeded with 8-10 L4 hermaphrodites, which were allowed to develop into adults and lay eggs for 24 h at 20 °C before being removed. The progeny were then grown for a further 48 h at 20 °C to the young adult stage.

Young adults were washed from the plates with M9 buffer and collected on a 40-µm cell strainer. Worms were collected into 1.5 ml Eppendorf tubes and washed three times. For each wash, they were pelleted by centrifugation at 600 × g for 6 min at 22 °C, the supernatant was aspirated, and the worms were resuspended in 1 ml M9 containing 0.1% PEG. Embryos were then isolated by bleaching the gravid adults. Bleaching solution (1.8 ml) was prepared immediately before use by mixing 750 µl H₂O, 600 µl 5 M NaOH and 450 µl 5% sodium hypochlorite (Thermo Fisher Scientific, 419550010). The worm suspension was distributed between tubes so that each contained approximately 10 µl of visible worm pellet. Each tube received 450 µl M9 with 0.1% PEG and 150 µl bleaching solution, and was shaken at 1,400 rpm and 22 °C for 13 min on a benchtop thermomixer. Progress of bleaching was observed by light microscopy. Immediately afterwards, the embryos were pelleted by centrifugation at 600 × g for 6 min at 22 °C, the supernatant was removed, and the embryos were resuspended in M9 with 0.1% PEG. This process was then repeated twice to ensure removal of bleaching solution. At the end of the third centrifugation, embryos were resuspended in approximately 100 μL of 20 % Ficoll in M9 as a cryoprotectant and immediately taken for freezing.

### Formvar coating of EM grids

Molybdenum 100-mesh EM grids without support foil (Plano, G2462MO or Graticules Optics, PY100-MO) were coated with Formvar to obtain a support film approximately 70-150 nm thick. Prior to coating, grids were cleaned with acetone and allowed to dry. Clean, scratch-free glass slides were immersed for 10 s in a 1% (w/v) Formvar solution in chloroform in a glass funnel equipped with a tap, after which the solution was drained to form a thin Formvar film on the slide. The coated slides were allowed to dry at an angle for 3-5 min. The Formvar film was cut into a rectangle on both sides of the slide using a razor blade. The slide was then slowly lowered into a water-filled glass container, allowing the Formvar film to detach and float on the water surface. Clean grids were placed onto the floating film in four rows. The film carrying the grids was subsequently transferred onto a clean glass slide equipped with an absorbent paper sticker by rapidly bringing the edge of the slide into contact with the floating film. The coated grids were allowed to dry on the slide for approximately 24 h before use.

### High pressure freezing using the waffle method

Formvar coated molybdenum 100-mesh grids were glow-discharged for 90 s on each side using an EM ACE200 (Lecia Microsystems GmBH) system at 15 mA and placed on palmityl-palmitate-coated 6mm Au/Cu planchettes (Wholwend Engineering)^7^. A 2 µL aliquot of freshly isolated *C. elegans* embryos was applied on top of the grid, which was then covered with a second planchette and immediately high-pressure frozen using a Leica EM ICE (Lecia Microsystems GmBH). Following freezing, the grids were recovered from the metal planchettes. During this process a small hole was pierced through the ice of each grid close to the grid border to facilitate subsequent correlation by providing a reference point visible in both light and electron microscopy. Grids were then clipped into cartridges suitable for FIB milling, and stored in liquid nitrogen until further processing.

### Cryo-light microscopy

All grids were screened by light microscopy prior to FIB milling. Before mounting on the microscope, grids with excessive surface ice contamination were cleaned to remove surface ice by gently rubbing them against a cotton bud cooled in liquid nitrogen. Grids were loaded onto a cryo-correlative stage (CMS196v³, Linkam) under liquid nitrogen vapor and mounted on an LSM 900 Airyscan microscope (ZEISS). An overview of each grid was first acquired using a 5× objective (NA 0.2), followed by acquisition of a ∼50 µm Z-stack using a 10× objective (NA 0.45) in confocal mode with the appropriate fluorescence channels. These overview images were used to identify grid regions containing early embryos. Selected regions were subsequently imaged at higher magnification using either bright-field or confocal microscopy. Z-stacks spanning ∼25 µm with a 0.5–1 µm axial step size were acquired using a 100× objective (NA 0.75).

### Low magnification grid correlation

A low magnification 5x bright-field overview of the grid was imported into MAPS software (ThermoFisher Scientific) and correlated with an SEM atlas image of the grid. The hole pierced near the edge of the grid during sample preparation was used as one fiducial marker, together with a second recognizable feature, such as ice contamination, the center marker of the grid, or an air bubble. When no other suitable feature was available, the opposite edge of the pierced hole was used as the second fiducial. The 10× fluorescence Z-stack was subsequently imported and correlated with the 5× overview. ICY^47^ was used for figure generation.

### High magnification grid correlation

The 100x confocal or bright-field Z-stacks were imported into MAPS and initially registered to the previously correlated 5x and 10x images based on the positions of grid bars and the orientation of individual grid squares. The registration of each high-magnification Z-stack was subsequently refined against the corresponding high-resolution SEM tile set. Distinctive ice contamination features located on the grid bars served as natural fiducial markers for precise correlation. ICY^47^ was used for figure generation.

### Iterative correlative FIB-milling

All steps were performed in an Aquilos cryo-FIB/SEM (Thermo Fisher Scientific) system. After the initial correlation described above, grids were coated with GIS for 90 s and sputter-coated at 10 Pa and 1 kV for 30 s. Trenches were milled at 16 nA perpendicular to the grid surface (i.e. at a milling angle of 90°) around a 50 × 50 µm region centred on the ROI. The stage was then returned to the mapping position (35° stage tilt with a 35° iFLM shuttle), and all material more than 20 µm below the top of the slab was removed at 7 nA. The sample was then tilted in increments towards the final milling angle (10° relative to the ion beam, corresponding to a stage tilt of 7°), with material removed at each step and the current reduced to 3 nA as the final thickness was approached. After completion of the undercut, the slab was typically 10-15 µm thick. The top 1 µm of the slab, containing the highly reflective GIS layer, was then removed to enable iFLM imaging. A full iFLM stack of the slab was acquired at 2 µm axial steps and used to refine the position of the ROI. The slab was then reduced from 50 × 50 µm to 50 × 20 µm by milling perpendicular to the grid at 500 pA.

To allow smooth and consistent GIS deposition on the front of the slab, the front 2 µm of material was removed using a cleaning cross-section (CCS) pattern at 500 pA. An additional 90 s of GIS was then applied with the stage at 45.5° rotation and 20° tilt, which oriented the GIS needle perpendicular to the slab front. These stage positions may need to be adjusted on other microscopes, as needle positions vary. The top 1 µm of material, containing the GIS layer on the upper surface, was again removed to allow iFLM imaging.

For iterative fluorescence correlation, an iFLM stack of the slab was acquired with a Z step size of 0.5 µm, and the target depth was calculated in a Fiji script as follows. The stack was resliced into a series of YZ planes, which were rotated about the X axis by the milling angle (10°) so that the slab surface was horizontal. The top of the slab was identified as the upper edge of the brightest region in the reflection channel, corresponding to the GIS layer. To determine the center of the ROI, a line scan normal to the top of the slab was drawn through the fluorescent target. The center was taken as the midpoint between the two half-maximum points of the subsequent intensity profile. The depth of the target was calculated as the difference in Z between the GIS surface reference and the centre of the ROI, multiplied by 1.31 to account for refractive-index mismatch.

Once the ROI depth had been determined, material was removed iteratively at a milling angle of either 10° or 13°. After each round of milling, a new iFLM stack was acquired and the ROI depth recalculated. Repeating these steps compensated for stage drift, measurement inaccuracies and lamella deformation. Typically, two to three measurements were sufficient to thin the slab from 15 µm to 3 µm, and a further two to thin it from 3 µm to 1 µm. Below 3 µm, the iFLM Z step size was reduced to 0.35 µm, and stress-relief cuts were milled at 500 pA at a thickness of 3 µm. Below 1 µm, the rapidly diminishing fluorescence signal hampered further iterations. Milling currents were 3 nA at thicknesses above 4 µm, 1 nA between 4 and 2 µm, and 500 pA between 2 and 1 µm. Polishing was carried out at 100 pA, with the milling angle increased to 10.5° or 13.5° to achieve more even milling.

After polishing, grids were sputter-coated again at 10 Pa and 1 kV for 5 s to reduce charging of the final lamellae. The grids were then removed from the microscope and stored as quickly as possible in standard grid boxes with the lids firmly closed, to minimize ice contamination.

**Table 2:** Milling parameters and estimated times for iterative fluorescence correlation.

|  | Ion beam current | Stage tilt angle | Stage rotation | Pattern type | Pattern XY dimensions | Pattern Z depth | Estimated milling time per step per 1 lamella |
|---|---|---|---|---|---|---|---|
| Surface cleaning | 0.5 nA | 6-15° | -70° | None | None | None | For the entire grid 5-10 min |
| Bulk trench milling | 16 nA | 17° | 110° | CCS | 2 x 50 µm by 50 µm | 20 µm | 45 min |
| Undercut | 3-7 nA | 35-7° | -70° | CCS | 50 µm x 10 µm, with y reduced at higher tilt angles | 10 µm | 30-45 min |
| Surface GIS Removal | 0.5 nA | 2-7° | -70° | Rectangle | 50 µm x 1 µm | 5 µm | 5 min |
| Slab shrinkage | 3 nA | 17° | 110° | CCS | 50 µm x 15 µm | 5 µm | 20 min |
| Front edge cleaning | 0.5 nA | 0° | 110° | CCS | 50 µm x 2 µm | 5 µm | 3 min |
| Surface GIS Removal | 0.5 nA | 2-7° | -70° | Rectangle | 20 µm x 1 µm | 5 µm | 2 min |
| Rough 1 | 3 nA | 7° | -70° | Top and bottom rectangles | ~ 20 µm in X, ~10 µm distance between the patterns | Visual estimate | 3-10 min |
| Rough 2 | 1 nA | 7° | -70° | Top and bottom rectangles | ~ 18 µm in X, ~5 µm distance between the patterns | Visual estimate | 3-10 min |
| Rough 3 | 0.5 nA | 7° | -70° | Top and bottom rectangles | ~ 15 µm in X, 3 µm distance between the patterns | Visual estimate | 3-10 min |
| Stress relief cut | 0.5 nA | 7° | -70° | Line | 2x 3 µm lines opposed at 60°, forming a tip away from the end of the slab | Visual estimate | 1 min |
| Medium 1 | 0.5 nA | 7° | -70° | Top and bottom rectangles | ~ 15 µm in X, 2 µm distance between the patterns | Visual estimate | 3-8 min |
| Medium | 0.3 nA | 7° | -70° | Top and bottom rectangles | ~ 15 µm in X, 1 µm distance between the patterns | Visual estimate | 3-6 min |
| Fine | 100 pA | 7° | -70° | Top and bottom rectangles | ~ 15 µm in X, 500 nm distance between the patterns | Visual estimate | 2-5 min |
| Final Polishing | 49 pA | 7.5° | -70° | Top rectangle | ~ 15 µm x 200 nm, on the very top of the unmilled material | Visual estimate | 20-60 s |

### Bidirectional fluorescence-guided SOLIST

Serial cryo-FIB lift-out was performed using the SOLIST workflow combined with three-dimensional fluorescence-based targeting. A metal block approximately 15-20 µm in lateral dimensions was prepared from either a gold receiver grid (200 mesh UltrAuFoil R 2/2, Quantifoil) or a 200-mesh copper thin-bar grid (Plano) and attached to the micromanipulator needle by ion-beam-induced redeposition. Prior to attachment, the contact surface of the metal block was polished to provide a flat interface for attachment to the biological specimen.

Prior to milling, surface ice contamination was removed using a 0.5 nA ion beam current at a milling angle of 10°. Regions of interest identified by prior cryo-fluorescence microscopy were mapped onto the FIB/SEM data using MAPS software as described above. The sample was sputter-coated at 30 mA for 30 s at 0.1 mbar, followed by 2 min of GIS deposition and a second identical sputter-coating step.

For preparation of the lift-out block, the grid was oriented perpendicular to the ion beam (17° stage tilt, 110° rotation). A small cross-shaped fiducial was milled approximately 10 µm from the fluorescence-defined target using a 30 pA ion beam. Trenches were subsequently milled around an approximately 30 × 18 µm block while retaining an approximately 6 µm bridge connecting the block to the surrounding material. Coarse trench milling was performed using cleaning cross-section (CCS) patterns at 7-16 nA, with the patterns directed toward the block. The upper surface of the resulting block was polished using a 1 nA CCS pattern to generate an approximately 20 µm-deep smooth contact surface for attachment to the micromanipulator. The needle was positioned such that the metal block contacted the polished upper surface of the biological block. The two blocks were joined by ion-beam-induced redeposition using multiple regular cross-section patterns with multi-pass 1 at 1 nA. A deposition depth of 5 µm was used for gold connection blocks and 10 µm for copper connection blocks. After confirming attachment, the remaining bridge connecting the biological block to the bulk specimen was severed using a rectangular pattern at 3 nA. The block was then gradually raised and retracted from the specimen using the micromanipulator. After successful lift out, while the lift-out block was still attached to the needle, the shuttle was removed from the microscope to a freshly-cooled loading station, to allow loading of a 200 mesh UltrAuFoil (Quantifoil Micro Tools GmbH) receiver grid onto the microscope. The receiver grid was kept clean and free of contamination by minimizing its time in liquid nitrogen.

Lifted-out material was deposited onto clean regions of the receiver grid. No sputter coating was carried out to facilitate electrostatic attachment of the deposited slices. The stage was positioned at a milling angle of 6°, and the block was lowered until it made contact with the support foil of the selected grid square. Serial slices of approximately 4 µm thickness were released using a sequential milling pattern consisting of a non-milling reference region (marking the size of the future slice), a 0.5 µm wide CCS pattern operated from left to right, followed by a line pattern drawn along the top edge of the CCS pattern, all operated at 1 nA. After release of each slice, the remaining block was immediately moved away from the deposited material to minimize reattachment. For the slice containing the cross-shaped fiducial, a 5 µm slice was prepared with the cross positioned approximately in its center.

Following deposition, slices were imaged using the iFLM. Z-stacks were acquired to include the focal plane of the receiver-grid foil, enabling features visible in reflection images (i.e. holes in the grid foil) to be used for subsequent correlation. A new SEM tile set was acquired following iFLM imaging to account for stage displacement associated with rotation into the fluorescence-imaging position. The iFLM and SEM datasets were correlated as described above. The position of the fluorescence-defined target within each deposited slice was then used to determine the region retained for subsequent lamella preparation. Deposited slices were typically trimmed to an approximately 14 × 14 µm region, from which a final lamella approximately 10 µm wide and 14 µm long was prepared.

Prior to thinning, the front and back surfaces of each deposited slice were coated by sequential sputtering and GIS deposition. Samples were sputter-coated for 30 s at 30 mA and 0.1 mbar. GIS deposition was performed from opposing orientations for 15 s each to coat the top, front, and back surfaces of the slices. The back of each slice was then trimmed at 3 nA, while the front was initially milled at 3 nA and subsequently polished using a 1 nA CCS pattern. The samples were subsequently sputter-coated again, followed by two 45 s GIS depositions performed at different stage rotations to obtain a uniform protective layer on the front surface. After an additional sputter-coating step, an SEM image of the cleaned slices was registered with the corresponding fluorescence data in MAPS. The final lamella position was selected to retain the fluorescence-defined target near the center of the lamella. The required vertical offset between the target height and the lamella position was determined from the distance of the target from the cleaned front surface and the FIB milling angle using trigonometric correction.

Lamellae were milled at a 6° milling angle. Based on the fluorescence correlation and calculated geometric correction, a 200 nm non-milling reference pattern was positioned at the calculated height above the reference cross plane and used to define the center of the final lamella. Initial top and bottom milling patterns, each 14 µm wide, were positioned 4 µm apart and progressively brought closer together while reducing the ion-beam current. Milling was performed sequentially at pattern separations of 4 µm (1 nA), 2 µm (0.5 nA), 1 µm (0.3 nA), 0.8 µm (100 pA), 0.5 µm (100 pA), and finally 200 nm (49 pA). Pattern positions were adjusted between milling steps when necessary to compensate for sample drift or beam-current-dependent shifts. Final polishing was performed from the back toward the GIS-protected front surface using a 0.5° overtilt and 30 pA current. Following lamella preparation, the grid was sputter-coated for 5 s at 7 mA and 0.1 mbar.

**Table 3:**
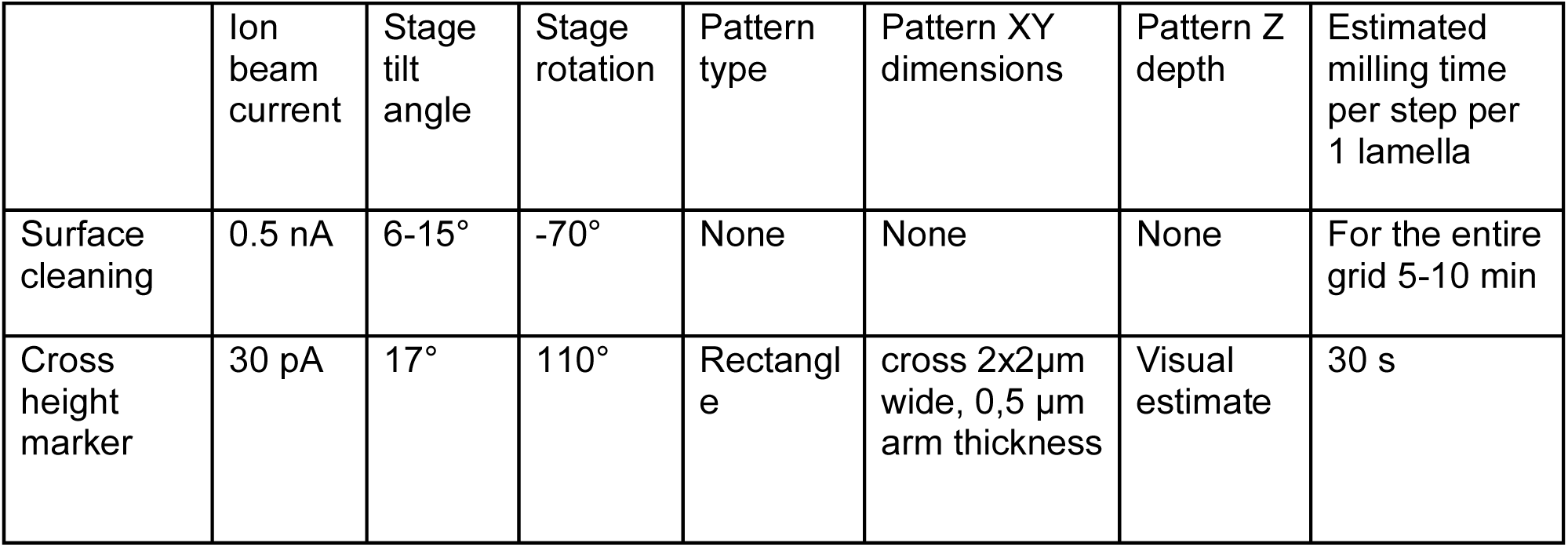
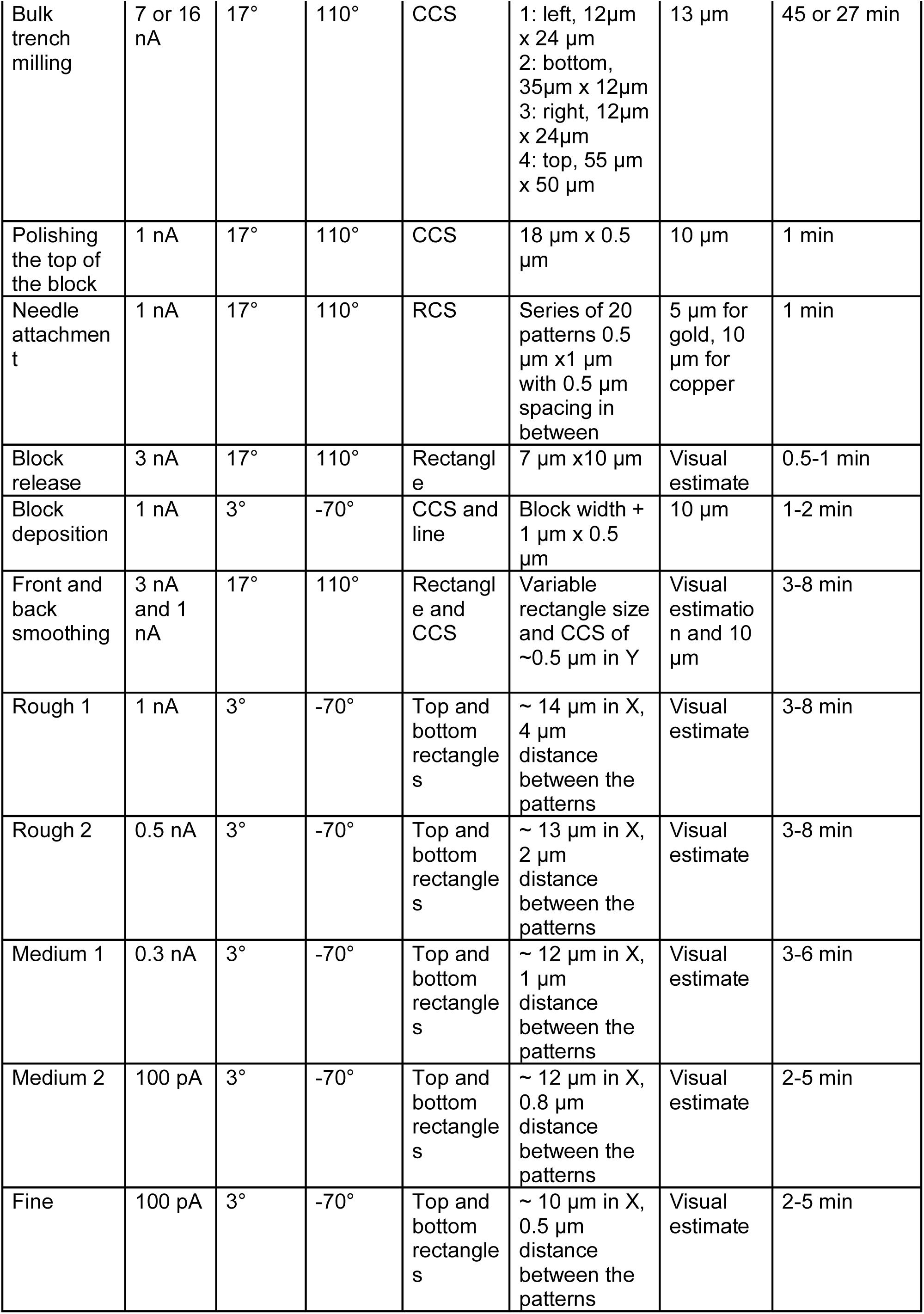
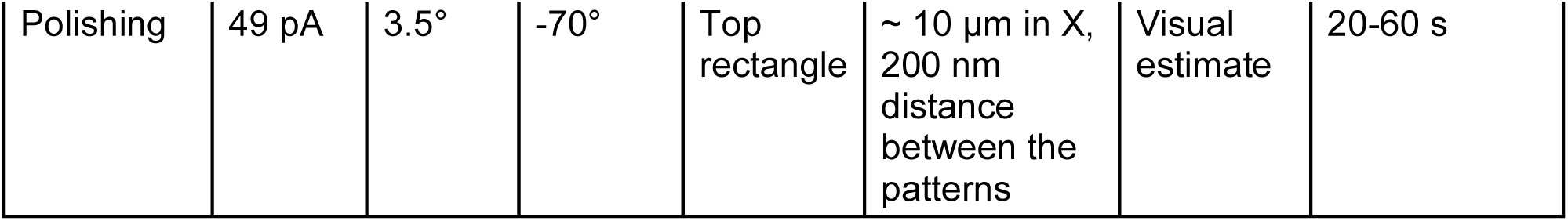
Milling parameters and estimated times for bidirectional fluorescence correlation.

### High-throughput lift-out and automated lamella preparation

The correlation steps followed the procedure described above, without the requirement for high precision. Trench milling was performed around the identified embryo with the waffle sample perpendicular to the ion beam (stage tilt 17° and rotation 110°). The preserved site was big enough to contain the whole embryo, around 50 by 35 µm. The top-side milled area was wide enough to accommodate the needle and copper block (40 µm in height), the sides and bottom patterns were 10 µm in width. The total trench milling time was around 20 minutes with 16nA CCS pattern and milling depth of 8 µm.

After the liftout of the block, the shuttle was unloaded to allow the insertion of an unsputtered receiver grid. After the receiver grid was loaded onto the stage, the liftout block was sliced onto the receiver grid with a 0.5 µm height CCS pattern, followed by a line pattern, leaving each slice 4 µm in thickness. A 50 µm height liftout block resulted in 10 slices. After the lift-out block is exhausted, a second lift-out and further slices deposition can be performed if the sample and time allows. iFLM images were taken and overlayed onto the SEM image of deposited slices in MAPS.

Two thin layers of 15s GIS were applied onto the slices, with T(stage tilt)=35° R(rotation)=- 70°, and T=-10° and R=110°, respectively. Afterwards, a sputter coating layer was applied to minimize charging.

The receiver grid was positioned perpendicular to the ion beam to cut the front leading edge and optionally the back far-edge with 3nA CCS depth of 2 µm. The position of the milling pattern was based on the fluorescence signal, so the ROI is ∼ 5 µm in the remaining slice and in the center. Then the stage was tilted to 0° and the front edge further smoothed with 1nA CCS 2 µm in depth.

An additional layer of 90s GIS was targeted to the fresh-cut surface with the grid positioned at milling angle and 2mm away from the coincidence point. Sputter coating was applied to reduce charging during automation.

AutoTEM cryo (Thermo Fisher Scientific) was used for automated lamellae preparation with a 10°milling angle. Fiducial features, such as star-shaped markers(with rectangle patterns 1×0.2×0.2um), were milled with 1nA into the surface of the slice to improve the accuracy of correlation between different ion-beam currents during automated milling (Supplementary Fig. 5) Rough milling was performed sequentially at pattern offset of 2 µm (1 nA), 1 µm (0.5 nA), 0.5 µm (0.3 nA). The thinning was performed with a pattern offset of 200nm (50 pA) and 0 nm (30 pA).

Following completion of the automated thinning, iFLM images on the thinned lamellae were taken to assess which lamellae still contained fluorescence signals from the targets. Then the lamellae were further polished manually from the top using a 0.5° overtilt to achieve the desired final thickness. Before the grid was removed from the microscope, a light sputter-coating step of 5 s at 7 mA and 0.1 mbar was performed to reduce charging during subsequent TEM imaging.

**Table 4:** Milling parameters and estimated times for high-throughput SOLIST.

|  | Ion beam current | Stage tilt angle | Stage rotation | Pattern type | Pattern XY dimension | Pattern Z depth | Estimated milling time per step per 1 lamella |
|---|---|---|---|---|---|---|---|
| Surface cleaning | 0.5 nA | 6-15 | -70° | None | None | None | For the entire grid 5-10 min |
| Bulk trenches milling | 16 nA | 17 | 110° | CCS | 1: left, 10 µm x 45 µm<br>2: bottom, 55 µm x 10 µm<br>3: right, 10 µm x 65 µm<br>4: top, 60 µm x 40 µm | 8 µm | 20 min |
| Polishing the top of the block | 1 nA | 17 | 110° | CCS | 18 µm x 0.5 µm | 10 µm | 1 min |
| Needle attachment | 1 nA | 17 | 110° | RCS | Series of 20 patterns 0.5 µm x1 µm with 0.5 µm spacing in between | 5 µm for gold, 10 µm for copper | 1 min |
| Block release | 3 nA | 17 | 110° | Rectangle | 3 µm x 20 µm | Visual estimate | 3 min |
| Block deposition | 1 nA | 4 | -70° | CCS and line | Block width + 1 µm x 0.5 µm | 10 µm | 1-2 min |
| Front removing | 3 nA | 17 | 110 | CCS | Variable size | 2 µm | 1-3 min |
| Front smoothing | 1nA | 0 | 110 | CCS | 40 µm x 1.5 µm | 2 µm | 2 min |
| Feature milling for autoTEM | 1nA | 7 | -70 | Rectangle | Series of patterns 1 µm x 0.2 µm | 0.2 µm | 1 min |

autoTEM parameters:

Milling angle: 10°, Lamellae size: 15μm x 5 μm, Correction factor 0.3, Final Thickness 200nm

**Table 5:** AutoTEM parameters used in automated milling for high-throughput SOLIST.

|  | Pattern Offset | Overtilt | Depth Correction | Milling Current | Pattern Overlap | Pattern Type |
|---|---|---|---|---|---|---|
| Rough milling | 2 µm | 0° | 350% | 1 nA | - | CCS |
| Medium Milling | 900nm | 0.5° | 160% | 0.5 nA | 400% | CCS |
| Fine Milling | 500 nm | 0.5° | 160% | 0.3 nA | 200% | CCS |
| Polishing 1 | 150 nm | 0° | 160% | 50 pA | 200% | CCS |
| Polishing 2 | 0 nm | 0° | 160% | 30 pA | 200% | CCS |

### Data Collection

For iterative fluorescence correlation, data were collected on a Titan Krios (Thermo Fisher Scientific) operated at 300 kV, equipped with a K3 camera (Gatan) and a Quanta GIF (Gatan) with a slit of 10 eV. SerialEM^48^ and PACEtomo^49^ were used to collect tilts between +50 and -70 degrees starting from a -10 degree pretilt, with 3 degree increments, for a total electron dose of 150 e-/Å^2^ over the whole tilt series. Defocus was set between -6 and -8 µm, with a pixel size of 2.094 Å.

For bidirectional fluorescence approach and high-throughput SOLIST methods, the data was collected on Titan Krios G4 transmission electron microscope (Thermo Fisher Scientific) operated at 300 kV, equipped with Falcon4i camera and a Selectris X energy filter with a slit width of 10 eV. Tilt-series acquisition positions were identified using SerialEM^48^ and collected with PACEtomo^49^. Images were acquired at a pixel size of 3.033 Å or 2.95 Å, with a target defocus ranging from −4 to −7 µm and an exposure time of approximately 1 s per tilt. Each tilt received 2.54 e⁻/Å², yielding a cumulative dose of approximately 140 e⁻/Å² per tilt series. Tilt series were collected using a dose-symmetric acquisition scheme, with 2° angular increments over a range of ±54° starting from the lamella pretilt.

### Tilt Processing & Tomogram Reconstruction

Frames were preprocessed using WarpTools^50^ for motion correction, CTF estimation, and frame averaging. Tilts were then aligned using Aretomo2^51^ and miss-alignment^52^, before being processed again and reconstructed using WarpTools. For denoising and missing wedge compensation, tomograms were processed using Isonet2^53^.

## ACKNOWLEDGMENTS

We thank Simone Köhler for sharing the labeled *C. elegans* line. We would like to thank Georg Wolff for help with formvar coating of EM grids, and for providing a waffle milling protocol. We acknowledge the access and services provided by the Imaging Centre at the European Molecular Biology Laboratory (EMBL IC), generously supported by the Boehringer Ingelheim Foundation. We acknowledge access to the cryo-EM infrastructure (HDcryoEM), support provided by the Heidelberg University cryo-EM platform, and access to the data storage service SDS@hd and high-performance computing cluster bwHPC-S5, which are funded and supported by the German Research Foundation (DFG), the Federal Ministry of Education and Research (BMBF) and the Ministry of Science Baden-Württemberg, among others, within the framework of the Excellence Strategy of the Federal and State Governments of Germany. This work was further funded by European Union’s Horizon Europe Project IMAGINE (GA#101094250). CP was supported by INST 35/1931-1 FUGG and the State of Baden-Württemberg through the Strukturund Innovationsfonds für die Forschung (SI-BW). F.T. and C.P. were supported through the DFG SFB/TRR 186/3-A03 (Project-ID 278001972) and the SFB 1638/1-Z03 (Project-ID 511488495).

## AUTHOR CONTRIBUTIONS

O.K., F.T. and S.M. conceptualized the work. A.B. prepared *C. elegans* cultures for HPF. A.B., O.K. and F.T. prepared waffle-frozen samples. O.K. and F.T. screened samples at cryo-LM. O.K., F.T. and Z.Y. performed FIB-milling experiments and collected cryo-ET data. O.K. and F.T. analysed the data. O.K. and F.T. wrote the initial draft. Final review and editing was done by O.K., F.T., Z.Y., C.P., and S.M. S.M. and C.P. acquired funding.

## AI Statement

Claude (Anthropic) and ChatGPT (OpenAI) were used to assist with editing the manuscript text. All content was reviewed and revised by the authors, who take full responsibility for the final manuscript. The Fiji script was developed using Claude.

## Data Availability

The Fiji scripts and accompanying documentation for iterative fluorescence correlation are available in the EMBL GitHub repository (https://github.com/embl-ic/mattei-research/tree/main/iterative_fluorescence_correlation).

## Supplementary figures

**Supplementary figure 1.**
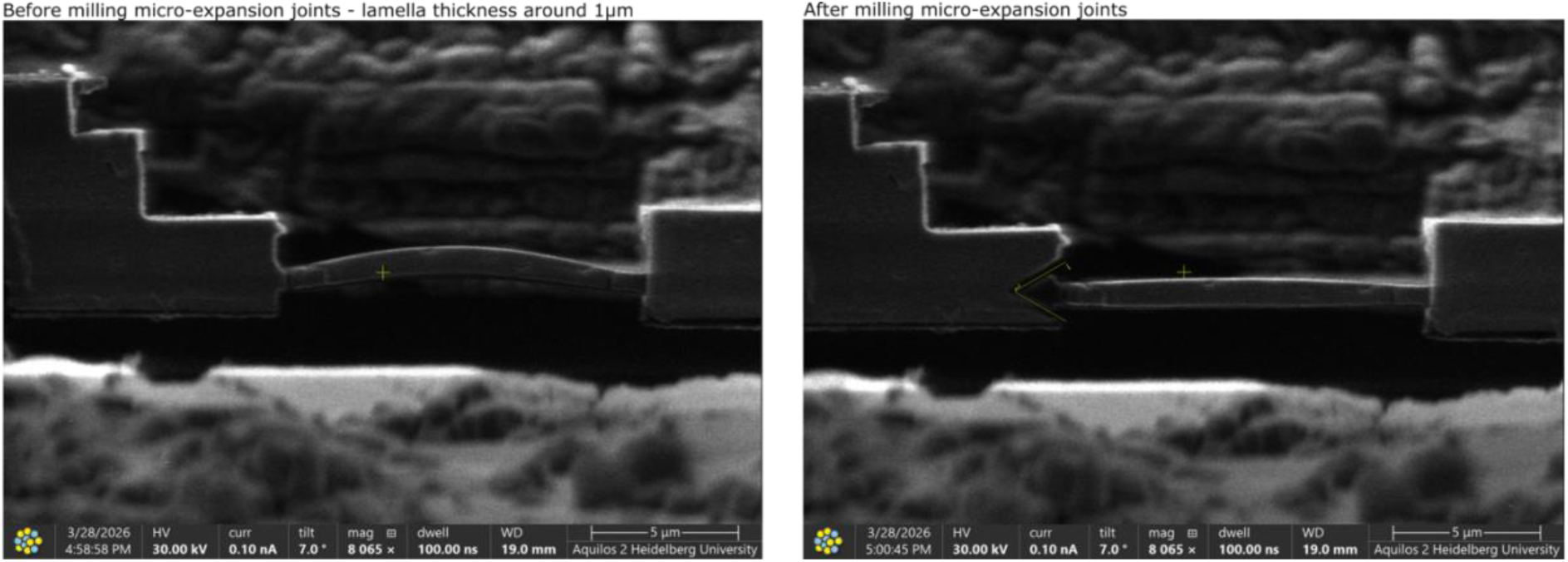
Stress-relief cuts reduce slab bending during iterative thinning. FIB images of the same approximately 1-µm-thick slab before (left) and after (right) milling a micro-expansion joint. The bowed slab becomes straighter after the stress-relief cut. Scale bars, 5 µm.

**Supplementary figure 2.**
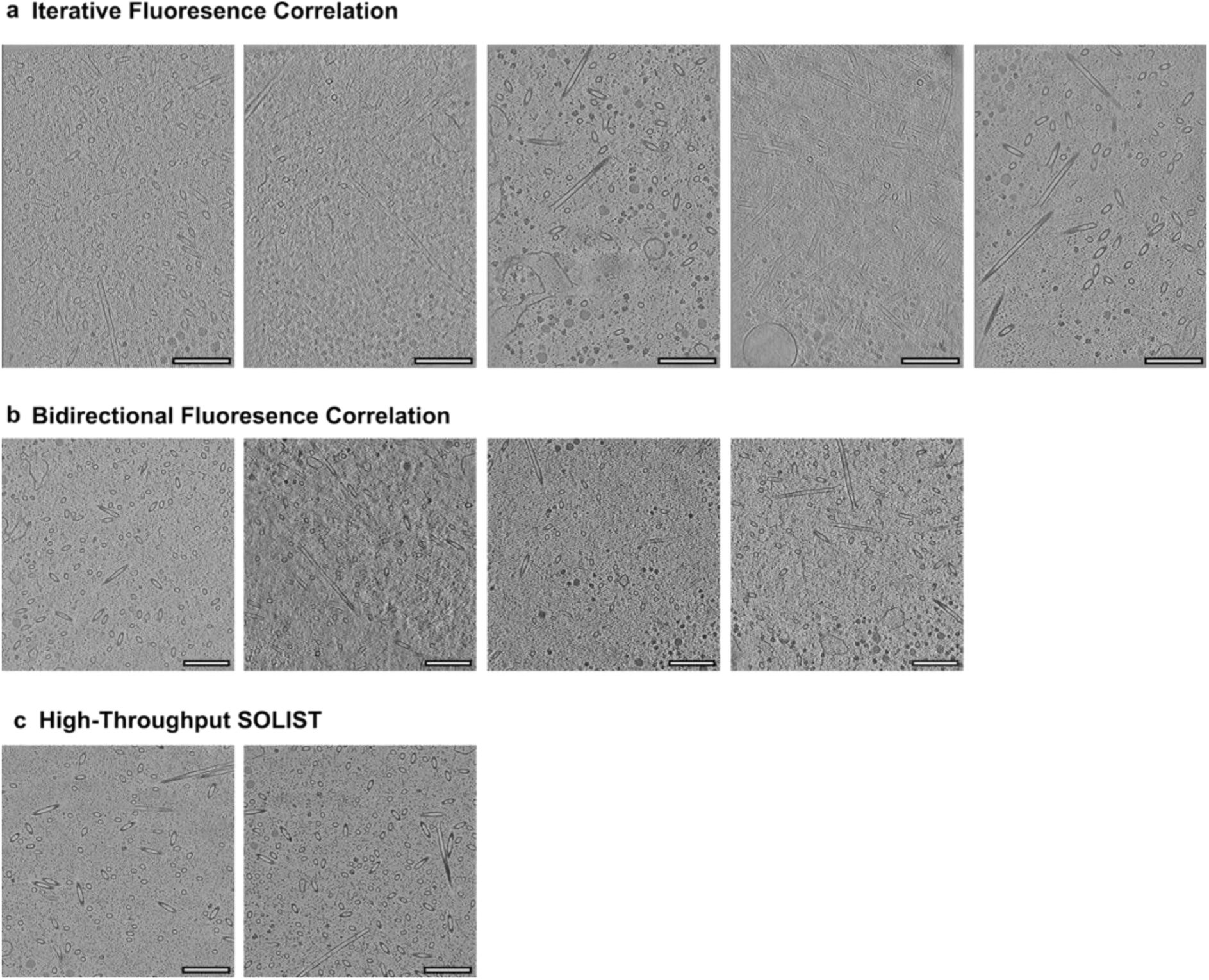
Centrosomes recovered using the three preparation workflows. Tomographic slices from target-positive lamellae prepared by iterative fluorescence correlation (**a**; five lamellae), bidirectional fluorescence correlation (**b**; four lamellae) and 2D-guided high-throughput SOLIST (**c**; two lamellae). Together, these images show the centrosomes recovered in all 11 target-positive lamellae reported in Table 1. Scale bars 200 nm.

**Supplementary figure 3.**
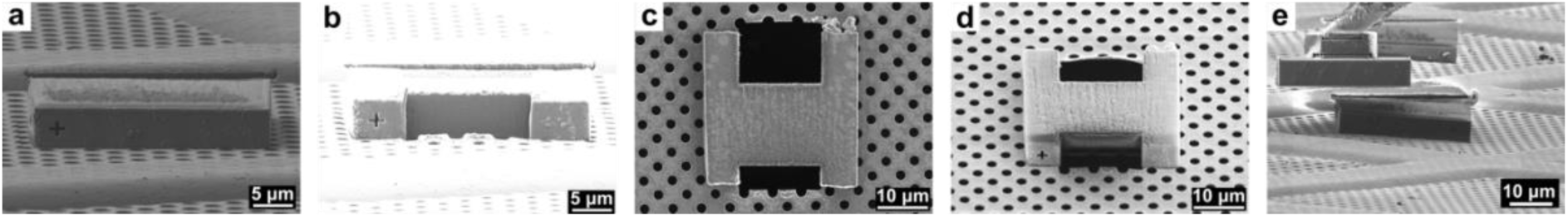
Support-foil flatness and trimming of deposited slabs for bidirectional targeting. **a**) FIB image showing how uneven support foil can alter the orientation of deposited slices. **b**) FIB image of a deposited slice before cleaning its front surface. **c**) The slice in b after front-surface cleaning. **d**) The same slice after removal of excess material behind the target. **e**) SEM image of the slice after front and back trimming. Scale bars, 10 µm (a,d,e) and 5 µm (b,c).

**Supplementary figure 4.**
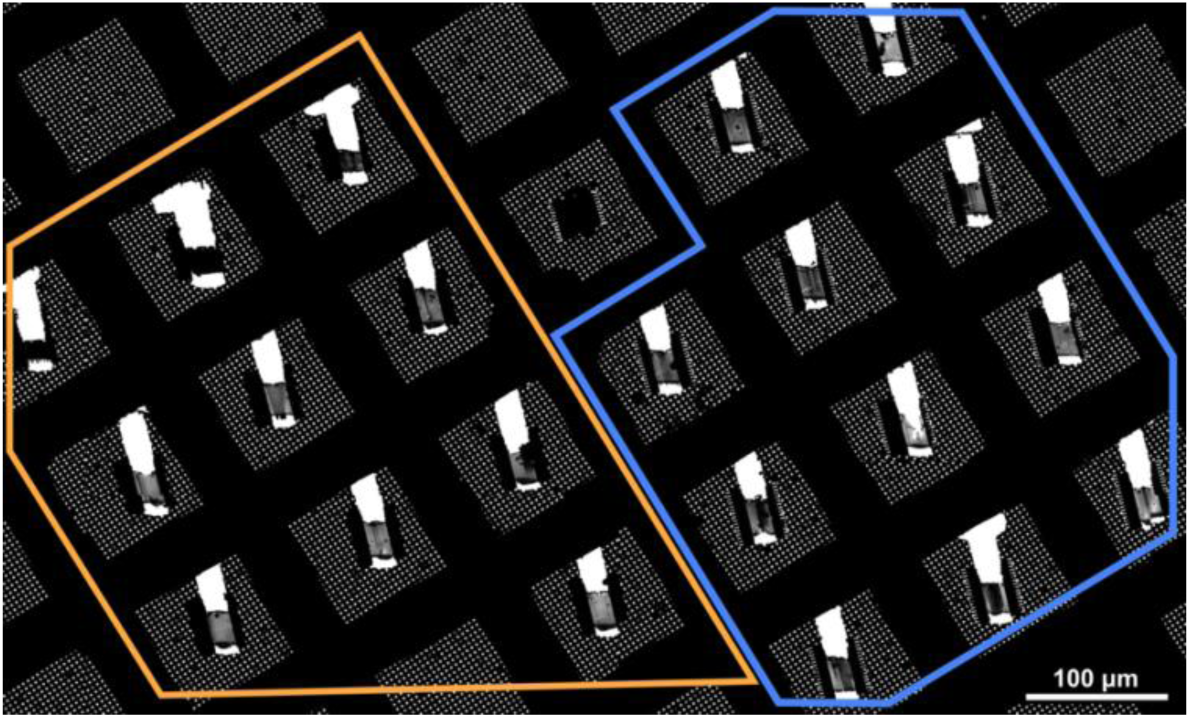
Survival of SOLIST lamellae prepared with and without micro-stitches on a receiver grid without sputter coating. TEM overview of a non-sputter-coated receiver grid after transfer. All 10 lamellae prepared with micro-stitches (orange polygon) and all 11 prepared without micro-stitches (blue polygon) remained attached. Scale bar, 100 µm.

**Supplementary figure 5.**
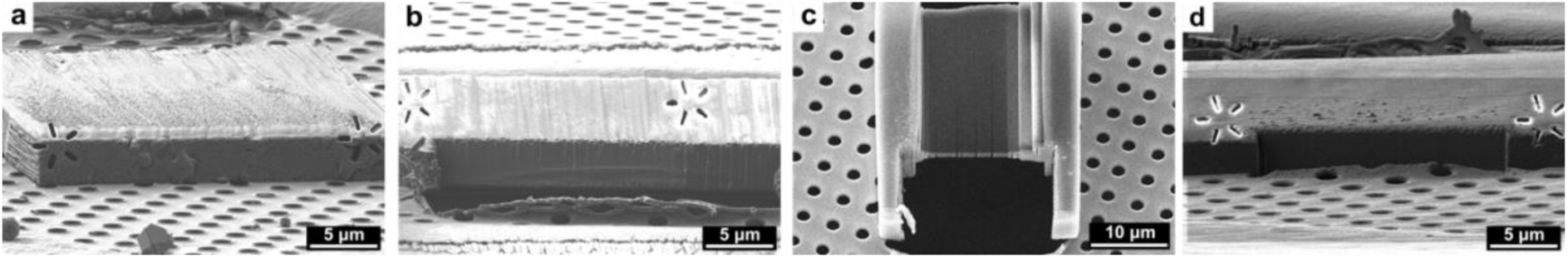
GIS coating and front shaping during SOLIST slice preparation. **a**) FIB image of a deposited slice before shaping its front surface. **b**) FIB image showing an irregular milling front beneath an uneven slice surface. **c**) SEM image of a lamella prepared from a slice with an uneven front, showing pronounced curtaining and disruption of the protective layer. **d**) FIB image of a slice coated using the gas injection system (GIS) from two directions before shaping, showing a more uniform milling front.

